# IMGT® Analysis of the Human IGH Locus: Unveiling Novel Polymorphisms and Copy Number Variations in Genome Assemblies from Diverse Ancestral Backgrounds

**DOI:** 10.1101/2025.07.18.665479

**Authors:** Ariadni Papadaki, Maria Georga, Joumana Jabado-Michaloud, Géraldine Folch, Guilhem Zeitoun, Patrice Duroux, Véronique Giudicelli, Sofia Kossida

## Abstract

Unraveling the genetic complexity of the human immunoglobulin heavy chain (IGH) locus provides valuable insights into the mechanisms underlying the efficacy and specificity of the adaptive immune response. Despite its crucial role, the IGH locus remains insufficiently characterized, with its allelic diversity and polymorphisms inadequately investigated. In this study, we present an analysis of the human IGH locus, incorporating 15 human genome assemblies from diverse ancestries, including African, European, Asian, Saudi, and mixed backgrounds. Through our examination of both maternal and paternal assemblies, we uncover novel IGH alleles, copy number variations (CNV), and polymorphisms, particularly within the variable (IGHV) region. Our findings reveal extensive and previously uncharacterized genetic variability in the constant (IGHC) region and distinct IMGT CNV forms across individuals. This research contributes to a significant enrichment of the IMGT® IGH reference directory, databases, tools and web resources and lays the groundwork for a comprehensive IMGT® haplotype database which can be progressively enriched to support future studies in population-specific immune profiles and adaptive immune related disease susceptibility, as comprehensive datasets become available. Such a resource promises to propel personalized immunogenomics forward, with exciting applications in cancer immunotherapy, COVID-19, and other immune-related diseases.

**GRAPHICAL ABSTRACT:** 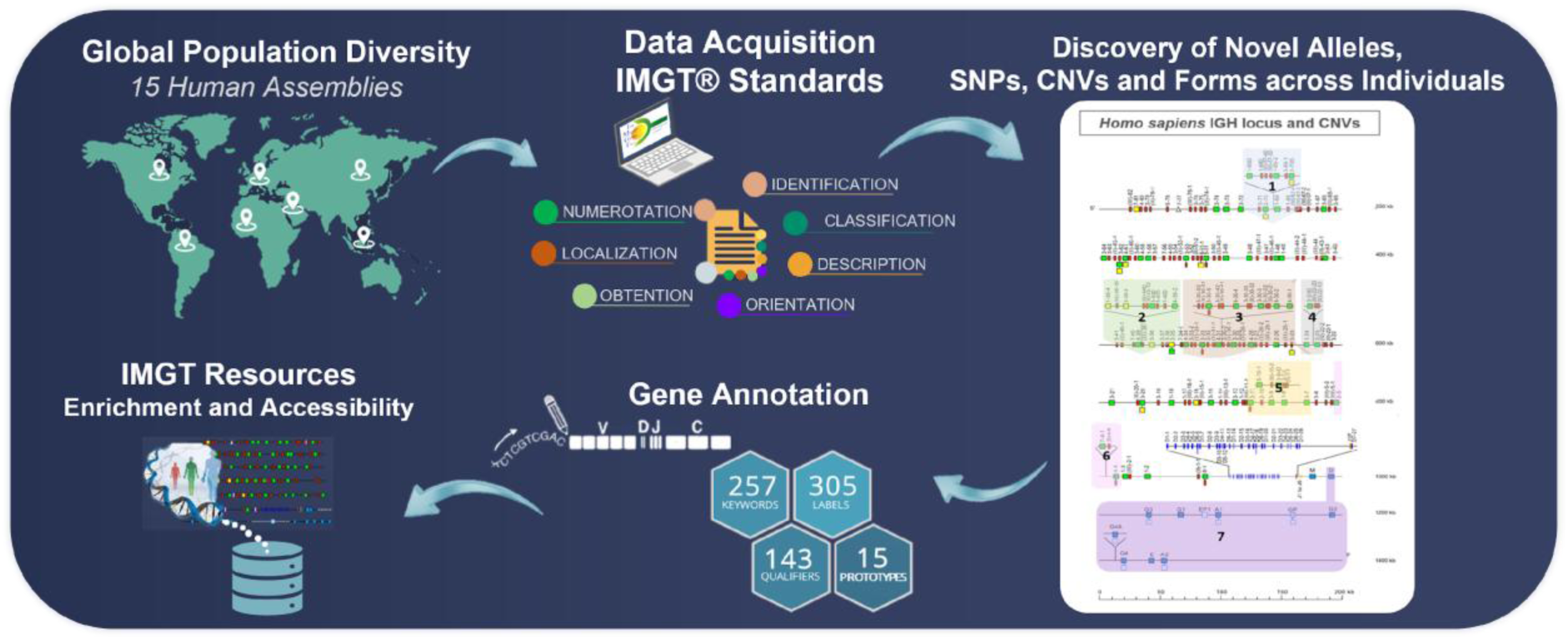

## INTRODUCTION

Located on chromosome 14q32, the human immunoglobulin heavy (IGH) locus is one of the most polymorphic and intricate, yet critical regions of the human genome. The IGH locus plays a central role in the adaptive immune response, orchestrating the production of the heavy chains of immunoglobulins (IG), which are crucial for antibody functionality (1). IGs or antibodies form an essential component to the immune system, as they recognize and neutralize a wide spectrum of pathogens, through highly specific antigen-binding sites (2,3). As such, the IGH locus not only supports immune diversity but also provides insights into how genetic variations may influence an individual’s resistance or vulnerability to diseases, positioning it as a key area of study in immunology and personalized medicine (4).

The remarkable diversity of IG repertoires, unique to each individual, arises largely from the complexity of the IGH locus. This diversity is primarily generated through VDJ recombination, a somatic gene rearrangement process that meticulously assembles one of the multiple variable (IGHV), with, in most cases, one diversity (IGHD), and one joining (IGHJ) genes into a productive V-D-J-REGION potentially capable of antigen binding.

Subsequently, splicing processes the primary mRNA transcript to link the rearranged V-D-J-REGION to a constant (IGHC) gene, enabling the synthesis of a productive IGH chain, which once associated with an immunoglobulin light chain will give an IG with antigen specificity (5). Human antibodies are categorized into five distinct classes, IgM, IgD, IgG, IgA, and IgE, and divided into 9 subclasses, one for IgM, IgD and IgE, two for IgA (IgA1 and IgA2) and four for IgG (IgG1, IgG2, IgG3, and IgG4) with their specific properties and functions largely determined by the constant region encoded by their respective IGHC genes (6).

The human IGH locus is not only structurally elaborate, containing numerous IGHV, IGHD, IGHJ, and IGHC genes, but is also subject to genetic variations such as single-nucleotide polymorphisms (SNPs), insertions and deletions, copy number variations (CNVs), and IGHC gene allotypes (Supplementary Figure S1), which significantly shape inter-individual immune responses and can impact susceptibility to various diseases (7,8). These variations hold implications for immunotherapy development, as they can reveal potential genetic adaptations that influence vaccine efficacy and immunological responses across populations (3,9,10).

While the importance of the IGH locus in shaping the human immune defense is well established, comprehensive studies of this critical region, particularly those utilizing data from individuals across different ancestries, remain scarce. The complexity and extensive polymorphism of the IGH locus pose substantial challenges for comprehensive analyses, leaving significant gaps in our understanding of its variability across different human individuals and populations. Prior analyses on the IGH locus have often relied on fragmented genome segments and scaffolds, which have only begun to unveil the rich genetic landscape of the IGH locus (11). Current publicly available reference genomes, such as GRCh38 (12) and T2T-CHM13 (13), serve as valuable starting points but indicate that much more remains to be discovered; structural variants, regulatory elements that influence the gene expression, population-specific characteristics (14,15). Evolving genomic research promises to significantly advance our understanding of the genetic diversity within the IGH locus across individuals and populations, improving the precision of immunotherapies in the management of autoimmune diseases, cancers, and infectious diseases.

IMGT® (International ImMunoGeneTics Information System®, (http://www.imgt.org) (16) has emerged, since 1989, as a comprehensive resource for the investigation of IG gene variations across various species, including humans, non-human primates, experimental models, and other jawed vertebrates. By providing extensive databases and analytical tools for the exploration of the structure and function of IG genes, IMGT facilitates the study of IG loci diversity and its implications for immune responses in vertebrates (16). However, comprehensive analyses of the IGH locus from individuals of different ancestries have been hindered by the lack of a holistic reference set that captures the full spectrum of genetic variation, for IGHV, IGHD, IGHJ and particularly IGHC genes. This study contributes to address this gap by presenting a novel collection of IGH locus analyses from individual genomes across diverse geographic origins, significantly updating the IMGT reference set. Utilizing genomic tools and meticulous methodologies, we explore, identify, and characterize novel allelic polymorphisms, CNVs, and IMGT CNV forms, as well as provide a comprehensive analysis of constant genes, highlighting the allelic diversity of the IGH locus across publicly available genome assemblies from individuals with diverse ancestral backgrounds. Our findings provide critical insights into the complex dynamics of individual immune response variations and lays the foundation for the establishment of a haplotype reference, which will be essential for future population-scale immunogenomic studies and precision immunology.

By comprehensively mapping and annotating the full spectrum of IGH genetic diversity, we contribute to the ongoing research aimed at the engineering of antibodies precisely tailored to individual genetic profiles. This work supports the advancement of personalized antibody immunotherapeutic strategies, improving the precision of therapeutic interventions and potentially enhancing treatment outcomes across a wide range of immune-related conditions, including cancer, COVID-19, and other diseases influenced by immune system variability (17,18).

## MATERIAL AND METHODS

The IMGT biocuration pipeline, as previously described (19), was performed for the analysis of the new human genomic data. To achieve precise and consistent results, the pipeline includes a combination of manual detailed work, as well as internally developed software tools, including IMGT/LIGMotif (20), NtiToVald (21) and IMGT/Automat (21), which are based on the IMGT-ONTOLOGY axioms and concepts: “IDENTIFICATION”, “DESCRIPTION”, “CLASSIFICATION”, “NUMEROTATION”, “LOCALIZATION”, “ORIENTATION”, and “OBTENTION” (22).

### Quality Assessment and Validation of Genomic Assemblies

The recent increasing availability of human genomic data, encompassing complete chromosomes, scaffolds, and contigs, necessitates the application of stringent quality control measures to ensure the accuracy and reliability of the analysis. In our study, we employed a two-phase methodology to select and validate genomic assemblies (Figure 1).

**Figure 1.**
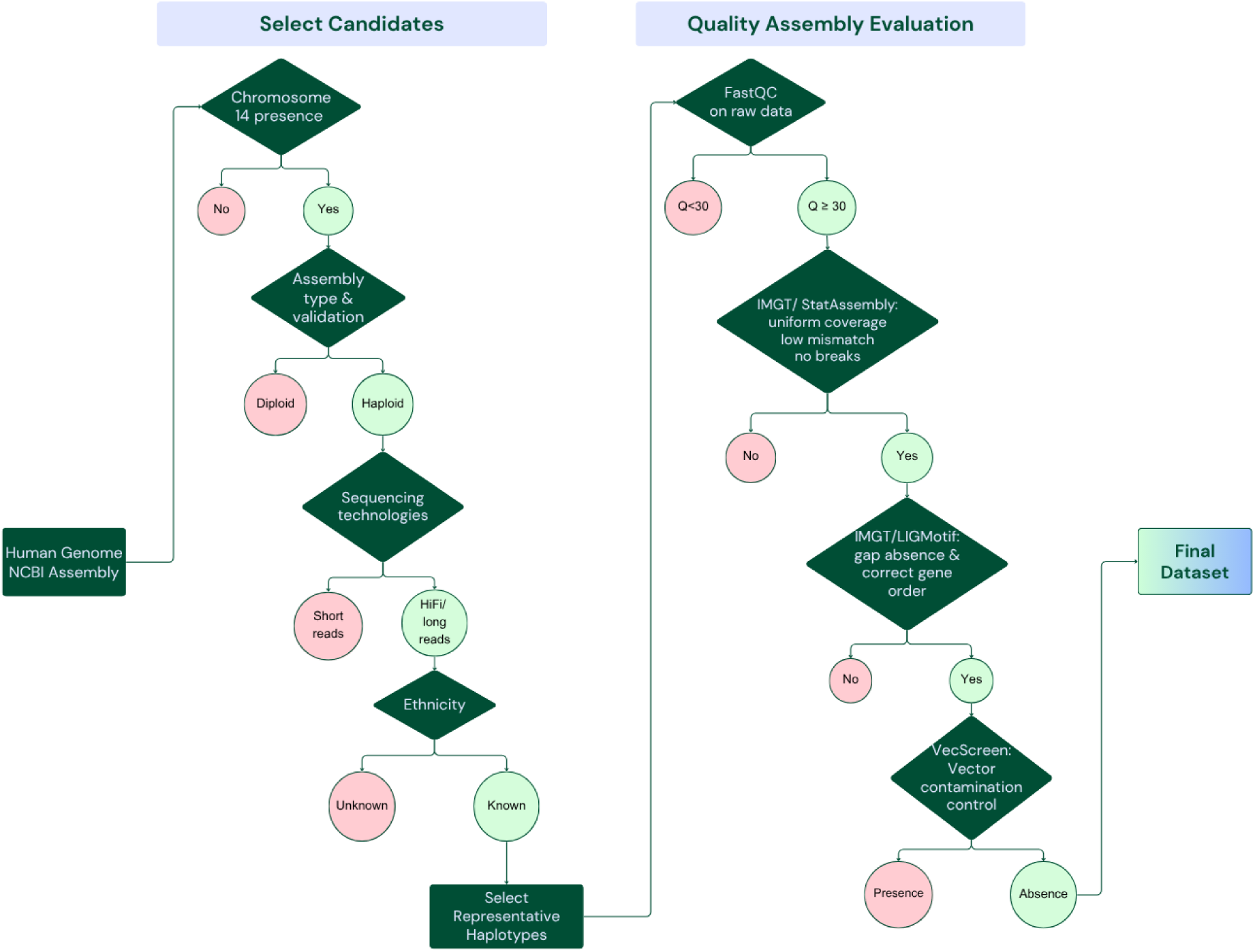
Decision pipeline for selecting high-quality human chromosomal assemblies. This flowchart illustrates the filtering steps applied to the available NCBI human assemblies to select the final dataset of this study. Assemblies were retained if they: (i) included chromosome 14, (ii) were haploid (validated through metadata and literature), (iii) were sequenced using long-read technologies (preferably HiFi), (iv) known donor ethnicity (excluding the reference assemblies T2T-CHM13v2.0 and GRCh37.p13). Then the selected assemblies had to pass stringent quality checks using FastQC (26), IMGT/StatAssembly (25), IMGT/LIGMotif (20) and VecScreen (29). Assemblies failing any step were excluded from analysis.

### Quality Assessment through NCBI Metadata, based on IMGT Pre-selection Criteria

All chromosomal assemblies available on NCBI (23,24) until December 1st, 2024, were evaluated to ensure complete coverage of the IGH locus (Supplementary Table S1). We followed the pre-selection criteria established by IMGT (available at: IMGT > Knowledge > IMGT Scientific Chart > IMGT rules for quality assessment of IG and TR loci in genome assemblies), which prioritize assemblies based on several key factors: assembly level and type, data recency, sequencing and assembling techniques, availability of associated raw sequencing reads, sequence gaps, and detailed sample information such as cell type, provider, literature validation, and ethnicity (https://www.ncbi.nlm.nih.gov/datasets/docs/v2/glossary).

Notably, chromosomal assemblies were prioritized, as they provide the most reliable, less fragmented and highly contiguous representation of an individual’s genome and ensure that the IGH locus is correctly mapped, on chromosome 14, telomeric extremity of long arm. The NCBI reference genomes, assembly GRCh38.p14 (GenBank Assembly ID: GCA_000001405), GRCh37p.13 (GCF_000001405) and assembly T2T-CHM13v2.0 (GenBank Assembly ID: GCA_009914755) were prioritized, as they are provided at Chromosome and Complete Genome assembly level, correspondingly, offering the highest resolution, accuracy and recency available for genomic analysis.

Beyond the chromosomal assemblies, additional scaffolds and contigs available through the NCBI databases whole-genome shotgun contigs (wgs) and nucleotide collection (nr/nt) were also examined, with the ultimate goal to address gaps in the literature, validate the existence of genes and alleles and contribute to a more comprehensive understanding of IGH variation across diverse human assemblies. An important parameter of our analysis was the assembly type to be haploid or haploid-with-alt-loci, avoiding the complexity and frequent inaccuracy of the two sets of chromosomes found in the diploids. Validation of haploid status was conducted through a comprehensive literature review when necessary.

### IMGT Quality Control of IGH Locus through Long-Read Analysis

To ensure further accuracy and reliability of the selected assemblies, IMGT developed IMGT/StatAssembly (version v0.1.4) (25), a dedicated quality control tool designed for the analysis of long-read sequencing data. The software is publicly available at: https://src.koda.cnrs.fr/imgt-igh/statassembly.

Long-read sequencing data for each genome assembly were retrieved from NCBI BioProjects. Preference was given to HiFi reads for their superior base-level accuracy. When unavailable, alternative long-read data were selected based on read length and quality. Read quality was evaluated using FastQC (26). Then raw data aligned with the assembly using platform-specific mapping strategies, including Minimap2 for long-read data (27). IMGT/StatAssembly then processes the resulting BAM alignments by specifically analysing the IGH locus and evaluating key quality metrics such as read coverage and sequence mismatches to detect potential inconsistencies (https://imgt.org/IMGTScientificChart/Assemblies/IMGTassemblyquality.php).

Regions with insufficient read support or low sequence identity were flagged for further evaluation. The results were summarized through graphical and tabular outputs, providing a comprehensive assessment of the assembly’s reliability.

This additional validation step, developed by IMGT, enhances confidence in the selected assemblies by reinforcing confidence in IGH locus analysis.

### Extraction, Quality Evaluation and Integration of the Locus in IMGT/LIGM-DB

The extraction of the locus from an NCBI assembly, which is the first step of the IMGT biocuration pipeline (19), was performed by comparing the assembly with the existing and most recent human IMGT reference directory sets, using the nucleotide basic local alignment search tool, BLAST (version BLAST+ 2.16.0) (28). The locus was extracted either complete from the chromosome, or from various contigs or scaffolds within the same assembly. For the IGH locus, according to the IMGT pipeline, the locus extraction is facilitated by the identification of the boundary 5’ and 3’ IG genes respectively, and the delimitation is defined by extending by 10kb each end of the locus, to include any potentially undetected genes.

The evaluation of the extracted locus was achieved primarily with the automatic tool IMGT/LIGMotif (20), which revealed the orientation of the locus on chromosome, the presence of IGHV, IGHD, IGHJ and IGHC genes, their number, position and orientation in the locus, as well as the possible presence of gaps. Regarding the orientation, when IMGT/LIGMotif indicated a reverse locus orientation, the corresponding reverse-complementary sequence was extracted from NCBI. An additional step in the locus evaluation was vector contamination control, using VecScreen (29), to identify any foreign DNA segments, including vector, linker, adapter, and primer regions. The default VecScreen scoring system, which categorizes matches as strong, moderate or weak based on BLAST similarity scores, was applied and no hits were found.

For the loci that all the locus quality criteria were fulfilled, IMGT/LIGM-DB (8) entries were created, with corresponding IMGT accession numbers.

### IMGT Gene Annotation and Functionality Determination

The standardized gene annotation was performed as described in the IMGT biocuration pipeline (19). The locus was first analyzed with the tool IMGT/LIGMotif (20), and the IGHV, IGHD, IGHJ and IGHC genes were semi-automatically annotated. The outcome was integrated, visualized, and verified in VectorNTI (v. 11.5.3, Thermo Fisher).

The identification of new genes and alleles in each assembly was based on the comparison with IMGT reference sets available at the time. After integrating the locus of each assembly, the previously annotated data served as a foundation for identifying novel genes and alleles in the studied assembly through a rapid localization process. This process involves aligning the studied locus with the existing coding regions, and if a 100% sequence identity on V, D, J-REGION or on C exons is observed, the gene is assigned the corresponding name and functionality based on the reference sequence. The rest of the elements were also checked to ensure their functionality and updated if needed.

The coding regions were analyzed with IMGT/V-QUEST and then subsequently verified, along with the rest IMGT standardized labels, through BLAST (5) and Multiple Sequence Alignment (MSA) methods, conducted with Clustal Omega tool/EMBL-EBI (30), MAFFT (31), and Kalign (3.3.1) (32) IMGT standardized labels were described for all genes in germline (IGHV, IGHD and IGHJ) and undefined (IGHC) configuration, according to the IMGT prototypes (available at https://www.imgt.org/IMGTScientificChart/SequenceDescription/prototypes.html).

The annotation rules outlined in the IMGT Scientific chart are based on the main concepts of the ‘DESCRIPTION’ axiom of the IMGT-ONTOLOGY (22) and allow the detailed characterization of the gene and thus, its functionality definition (functional (F), open reading frame (ORF) and pseudogene (P)), according to the ‘IDENTIFICATION’ axiom of IMGT-ONTOLOGY (22).

The annotated gene and allele sequences were integrated and are accessible through IMGT/GENE-DB (33) and the IMGT reference directory (available at: https://www.imgt.org/vquest/refseqh.html), following the IMGT unique numbering based on the ‘NUMEROTATION’ axiom (22), which ensures framework regions (FR-IMGT) and complementarity determining region (CDR-IMGT) delimitation consistency across IG.

The corresponding amino acid sequences are available in IMGT/3Dstructure-DB and IMGT/2Dstructure-DB (34). Once integrated, the sequences are utilized by the sequence analysis tools, including IMGT/V-QUEST (35), IMGT/HighV-QUEST (9), and IMGT/DomainGapAlign (36). All the annotated genomic data are available in corresponding IMGT repertoire pages: Locus representation, Locus description, Locus in genome the assembly, Locus gene order, Locus Borne, Gene tables, Potential germline repertoire, Protein displays, Alignments of alleles, Colliers de Perles, and germline [CDR1-IMGT.CDR2-IMGT.CDR3-IMGT] lengths (available at https://www.imgt.org/IMGTrepertoire/).

The annotated sequences were submitted to the NCBI Third Party Annotation (TPA) (37) and TPA accession numbers are progressively assigned to the non-rearranged IGH loci, replacing the original IMGT accession numbers (https://www.imgt.org/IMGTrepertoire/numacc.php).

### IMGT Nomenclature

The nomenclature of the IGH genes and alleles was based on the well-established IMGT nomenclature, in use since 1988. IMGT nomenclature, is described in “IMGT gene name nomenclature for IG and TR of human and other jawed vertebrates” (available at: https://www.imgt.org/IMGTScientificChart/Nomenclature/IMGTnomenclature.php) and follows the “CLASSIFICATION” axiom of the IMGT-ONTOLOGY. The gene and allele names share detailed information about the type of the gene, the allele and its position, and specifically, for IGHV genes: the subgroup or clan (for truncated pseudogenes) and position from 3’ to 5’ end of the locus, for IGHD and IGHJ genes: the sets and position from 5’ to 3’ end of the locus and for IGHC genes: the encoded class and subclass.

The new alleles were identified by comparison of their coding regions (V, D or J-REGION for IGHV, IGHD and IGHJ and exons for IGHC genes) with the IMGT reference directory sets. Based on their position in the locus, the alleles were designated with increasing two-digit numbers.

The genes that shared 100% coding region identity with an already characterized gene found in a different position of the locus, were named as duplicated genes (name of the “initial” gene, with addition of the letter “D”).

If a gene exhibited a lower identity percentage and was located between two previously characterized genes within the locus, it was considered a new gene and was named based on its position.

### Evaluation of Newly Identified Alleles for Accuracy and Reliability

As current experimental methodologies and computational tools may lead to unreliable sequences, especially for highly polymorphic loci (38), additional steps of control were applied to evaluate the new alleles, ensuring their reliability and distinguishing true genetic variants from potential sequencing errors.

IMGT/StatAssembly (version v0.1.4), was employed for the allele verification (25). The software analyzes the BAM alignment, generated by mapping HiFi reads to their corresponding assembly, and evaluates read support at the precise genomic position of each annotated allele, i.e., at the start and end positions of the allele based on its coordinates. Alleles were considered well-supported if their exonic regions were fully covered by reads with 100% identity, and each nucleotide was confirmed by more than 10 reads with a match rate exceeding 80%, providing strong evidence for their existence.

Subsequently, NCBI Magic-BLAST (39) was performed, aligning the newly identified alleles with the associated raw data provided by the NCBI Sequence Read Archive (SRA) repository. The parameters included a 100% identity score (perc_identity) and 40 threads (num_threads). While this method was applied to all assemblies, the computational demands and extensive processing time required limited the ability to obtain results, highlighting the need for more scalable approaches to validating assemblies on a broader scale.

In addition, countTags (40), which counts occurrences of a set of k-mer signatures in a set of FASTQ files, was employed to quantify the presence of 31bp kmers around the SNPs of the novel alleles, with the associated HiFi reads. This approach provided additional validation of specific polymorphisms, reducing the likelihood of errors due to sequencing.

Finally, Standard Nucleotide BLAST (BLASTn, version BLAST+ 2.16.0) was performed on uncertain genes and alleles against the NCBI nr/nt and wgs database (BLAST+ 2.13.0) to explore potential confirmation of their presence in existing genomic data.

Cross-referencing newly identified alleles with various databases provided additional evidence supporting their existence and authenticity. After careful consideration, alleles that did not meet the required validation criteria were not integrated, or, if previously integrated but later determined to not be biologically valid, were removed. In such cases, the names of the removed alleles are not reused in the future and the updates are stored at the IMGT/GENE-DB data updates page (available at https://www.imgt.org/IMGTgenedbdoc/dataupdates.html). This combined approach enabled the validation of new alleles, ensuring their accuracy and reducing the likelihood of sequencing errors.

It is worth noting that the new IGHV alleles were assigned only when they did not participate in DJ or VDJ rearrangements, to avoid misinterpreting the possible somatic hypermutations as germline variants.

### Copy Number Variations (CNVs), IMGT CNV forms characterization and IMGT haplotype Database Construction

Copy Number Variations (CNVs) significantly characterize the human IGH locus, involving structural changes such as deletions, insertions, and duplications that alter gene copy numbers, and are numbered from 5′ to 3′ in the locus (7). The CNVs and their IMGT CNV forms, which describe the variability of the gene number found within a specific CNV, were identified by IMGT/LIGMotif. However, given the complexity of the IGH locus, which contains numerous highly similar genes, the correct gene identification within the CNV regions was challenging, notably for the intricate CNV3. For such cases, additional parameters were taken into account, including the gene order within the locus and the relevant literature. For further verification, we run different MSA methods, including Clustal Omega tool/EMBL-EBI and MAFFT, with the most convenient and accurate for this analysis being Kalign, with default parameters (progressive alignment, identity matrix for nucleotide sequences, gap penalties of 10 and 0.1 for open and extension gaps, respectively, and parallelization for faster computation). This method has demonstrated high accuracy when aligning large and distantly related sets of sequences, making it a particularly effective method for encompassing entire CNV regions along with the untranslated regions (UTRs) of the genes (32) . We compared the CNV regions of the studied IGH loci, with the reference assembly T2T-CHM13v2, as this sequence integrates all known CNVs, with all known genes present within those CNVs (13,41). New IMGT CNV forms were identified and designated with Latin letters in alphabetical order. The distinctive IMGT CNV forms in each individual’s IGH locus contributed to the formation of unique IMGT haplotypes, resulting in the creation of an emerging IMGT haplotype database that may provide valuable insights into genetic profiles and ethnic backgrounds, and can be further enriched and confirmed through the analysis of additional assemblies from a larger number of individuals.

### IGHC genes Allotypes Identification and New Polymorphisms Characterization

The study of the IGHC gene variations, which includes the identification of existent allotypes (8) and novel IGHC gene polymorphisms, was facilitated by the data available in the IMGT repertoire pages: Alignments of alleles, which is based on the ’IMGT exon numbering’ concept, and Protein displays and Collier des Perles, which are based on the ’IMGT unique numbering’ concept, both belonging to the ’NUMEROTATION’ axiom of IMGT-ONTOLOGY.

## RESULTS

### NCBI Assembly Evaluation and Selection

In this study, we evaluated all 600 human chromosomal assemblies available on NCBI as of December 1st, 2024 to uncover novel variability and better characterize the composition and organization of the V, D, J, and C genes within the IGH locus The assembly selection process was conducted in two distinct stages to ensure the highest quality and reliability of the data for IGH locus analysis.

### Initial Assembly Selection Based on NCBI Metadata

All 600 chromosomal assemblies available until December 1st, 2024, underwent a thorough first review based on key metadata provided by NCBI. Notably, from August to December 2024, a large number of chromosomal assemblies (515) were uploaded simultaneously, and due to the high volume and limited representation of the relevant chromosome, they were not prioritized for this analysis. More specifically, those submitted in November by the Human Genome Structural Variation Consortium, despite being categorized under chromosomal genome assemblies, provide only contigs (e.g., GCA_964199205.1, GCA_964198295.1, GCA_964198215.1). Similarly, assemblies uploaded between August 28 and 31, 2024, by The Jackson Laboratory are listed as complete genomes but also contain only contigs (e.g., GCA_964213075.1, GCA_964212235.1, GCA_964212405.1). Additionally, most assemblies submitted in October and November 2024 by the UCSC Genomics Institute, lack a sequence for chromosome 14, even in cases when nearly all other chromosomes are present (e.g., GCA_044166515.1, GCA_018469685.2, GCA_018469675.2). Upon contact with NCBI, it was clarified that these assemblies, generated as part of the human HPRC project, include only chromosomes that are considered well-assembled, with most containing 7–20 chromosomes while the remaining sequences are in unplaced scaffolds. Although a few of these recently uploaded assemblies include chromosome 14, given the substantial influx of new data, priority was given on analyzing the assemblies submitted before August 2024. From these remaining 89 assemblies, 58 assemblies were excluded: 53 due to partiality, 3 due to suppression and 2 marked as diploid (Supplementary Table S1).

### IMGT Quality Criteria Evaluation

The remaining 31 assemblies were subjected to a more detailed evaluation using the internal IMGT- developed assembly quality tool, IMGT/StatAssembly (Supplementary Figure S2), and IMGT’s stringent quality criteria, as described in Methodology. This stage ensured that the most complete and high-quality assemblies were selected, leading to a final dataset of 9 chromosomal assemblies that met all required criteria: NCBI reference assembly T2T-CHM13v2.0, GRCh37.p13, ASM3717755v1 and ASM3717763v1 (maternal and paternal haploid assemblies from a Saudi individual), hg002v1.0.1.mat and hg002v1.0.1.pat (maternal and paternal haploid assemblies from an Eastern European Ashkenazic Jewish individual), mHomSap3.mat and mHomSap3.pat (maternal and paternal haploid assemblies from an individual of mixed ancestry: African, European, Native American) and PGP1v1 (haploid assembly from a North-Eastern European male).

As part of this second-stage evaluation, we also annotated specific scaffold assemblies for cases where chromosomal assemblies were either unavailable or did not meet the IMGT criteria. Notably, for isolates HG01243 (from Puerto Rican individual) and NA19240 (from African Yoruba individual), both diploid and their respective maternal and paternal haploid scaffold assemblies were fully annotated to ensure comprehensive genomic coverage. Additionally, the maternal and paternal assemblies of the Asian isolate HG005 were also included in our dataset for broader population representation, as other Asian chromosomal assemblies did not meet the required IMGT criteria. Moreover, supplementary scaffolds were included in the analysis to provide further validation of alleles, when necessary (eg. JADCYG010001029, JADEBP010005792). The final 15 assemblies investigated in this study are presented in Table 1 (further information available: https://www.imgt.org/IMGTrepertoire/index.php?section=LocusGenes&repertoire=locusAssembly&species=human&group=IGH).

**Table 1.**
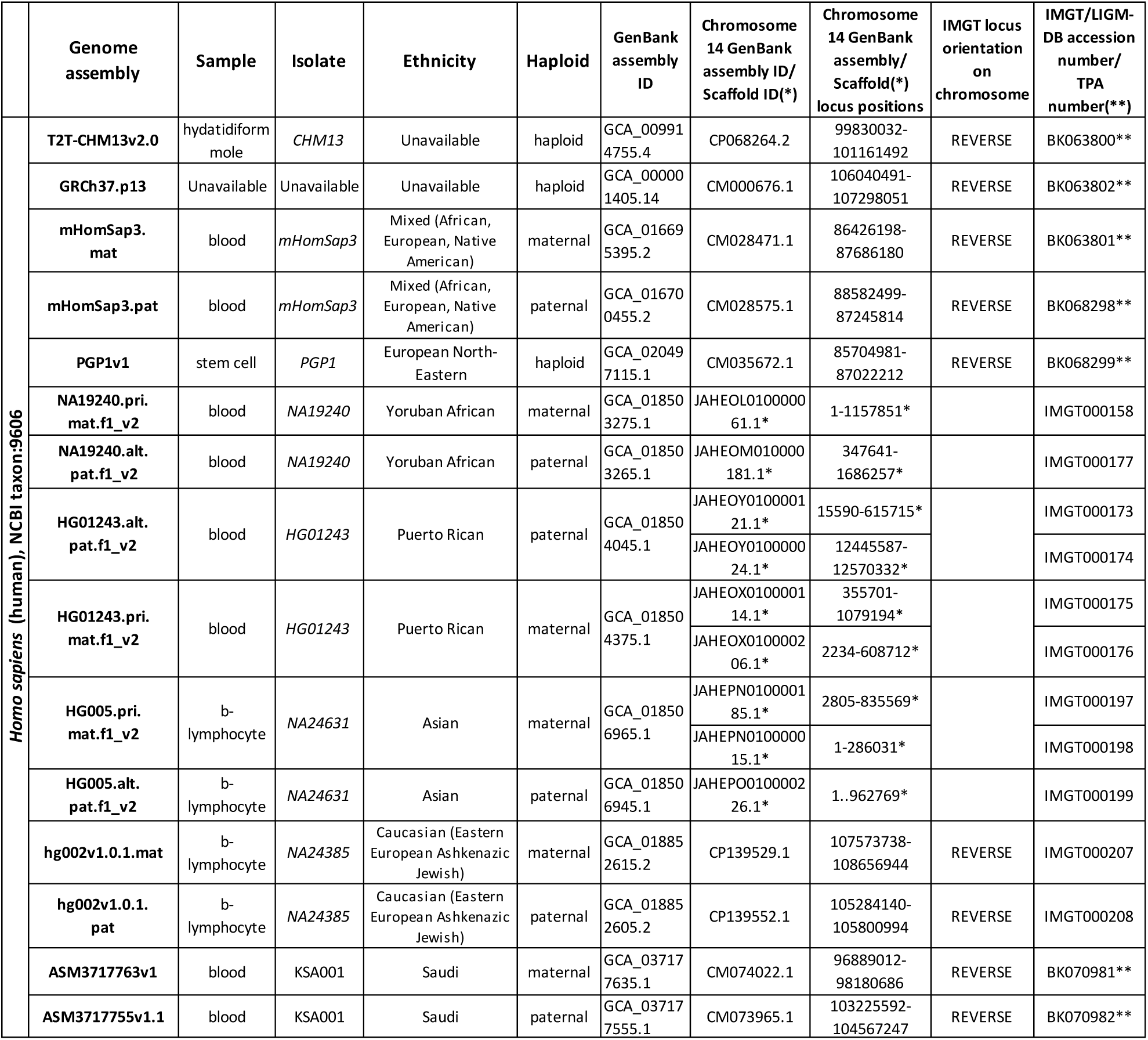
Information about genome assembly and IGH locus for 15 human annotated assemblies.

### 15 Diverse Genome Assemblies: Characterizing the IGH Locus Across Individuals of Different Ancestries

The IGH locus was extracted and annotated using IMGT® tools from 15 different genome assemblies publicly available at NCBI. These assemblies represent individuals with diverse ancestries, including African, Asian, European, Saudi, Puerto Rican, and mixed ancestries, along with two reference assemblies. The population information associated with each assembly in our study was carefully respected, as provided in the NCBI sample metadata, ensuring an accurate representation of the individuals’ genetic backgrounds.

The dataset comprised 8 assemblies encompassing the full germline IGH locus and 7 assemblies with rearranged loci. Notably, the deliberate inclusion of rearranged assemblies was pivotal, as it captures distinct stages of IG gene rearrangement occurring in B cells, reflecting the dynamic B cell maturation processes within the IGH locus. Also, three of the assemblies contained the complete IGH locus across two scaffolds.

Table 1 provides a comprehensive summary of the information of the selected assemblies, including sample origin, ethnicity, locus positions and corresponding accession numbers within the IMGT/LIGM-DB (42) and TPA databases (37). The reference assembly GRCh38.p14, already established in the IMGT databases, was used as a benchmark. The IMGT locus corresponding to this assembly is Homsap_IGH_1 and can be accessed here: https://www.imgt.org/IMGTrepertoire/index.php?section=LocusGenes&repertoire=locusAssembly&species=human&group=IGH. Also, the version, GRCh37.p13, was included in the dataset for updated consistency and reliability across analyses (BK063802).

To investigate the structural diversity and gene composition of the IGH locus across the assemblies studied, we analyzed key characteristics, such as number of genes and allele functionality (for information about novel allele functionality and distribution, see Supplementary Figure S3). Figure 2 provides an overview of the gene composition within complete, non-rearranged IGH loci across diverse human assemblies, highlighting the loci with the highest and lowest number of genes. The paternal assembly of the individual mHomSap3 of mixed ethnicity (BK068298, mHomsap.pat), had the largest IGH locus with 188 genes, including the highest number of IGHV genes (141). The reference assembly of the cell line CHM13 (BK063800, T2T-CHM13v2.0) and the paternal assembly of the Saudi individual KSA001 (BK070982, ASM3717755v1.1) contained the highest number of IGHC genes (12 each), due to the presence of the IGHG4 duplication (IGHG4A), while all the rest contained all the 11 previously described constant genes. The smallest IGH locus belongs to the maternal assembly of mHomSap3 (BK063801, mHomSap3.mat), with just 169 genes. Finally, despite variability in IGHV and IGHC gene numbers, the number of IGHD and IGHJ genes remained highly conserved across all assemblies, with 27 and 6 genes respectively, highlighting the lower structural variability of these regions (Figure 2).

**Figure 2.**
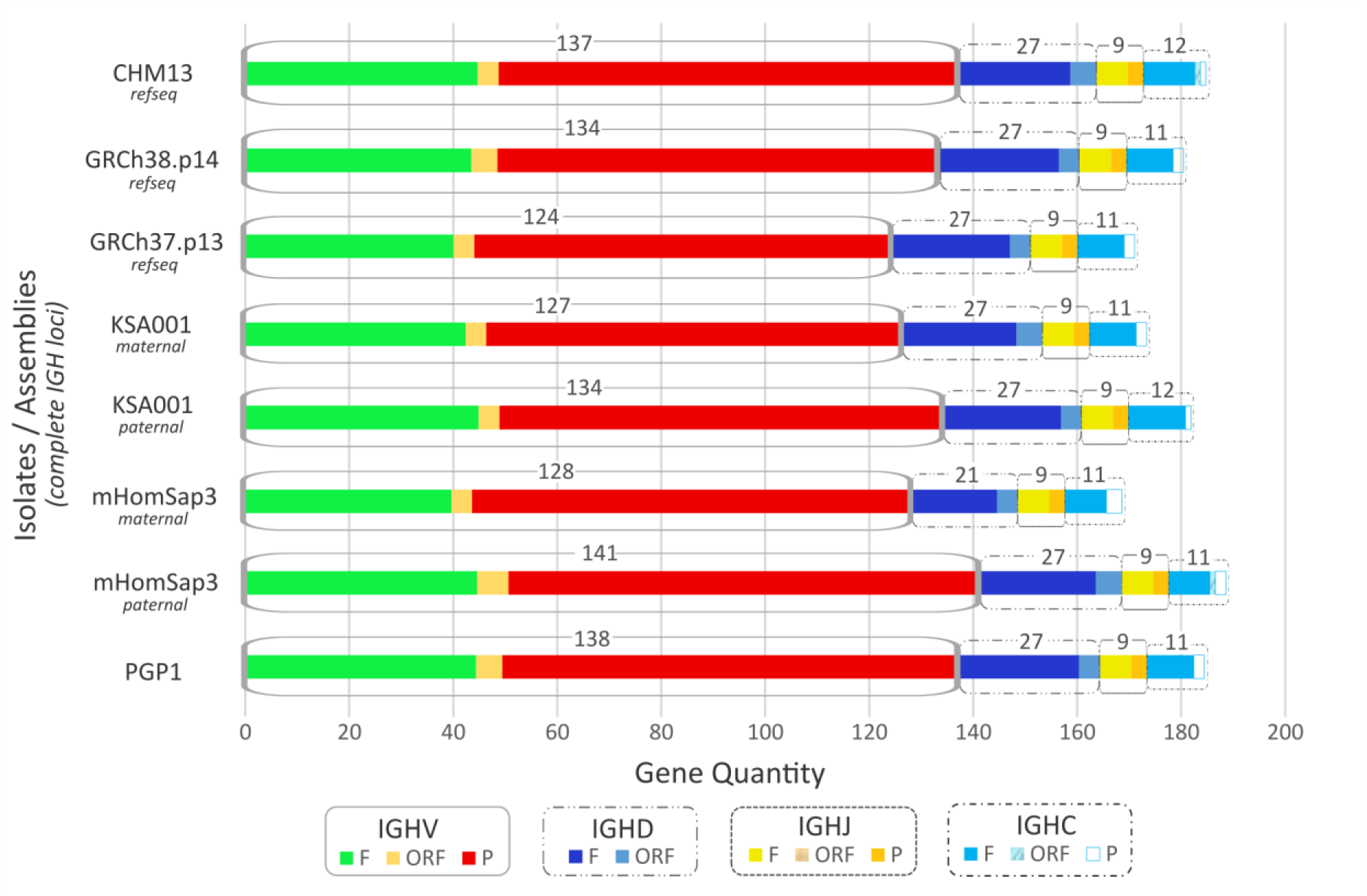
Gene composition and functionality classification of complete, non-rearranged IGH loci within 8 analyzed human assemblies. This graph displays the gene count and functionality distribution for each gene type (V, D, J, and C). Genes of the same gene type are grouped in brackets, with colors applied according to the IMGT color menu, reflecting the IMGT functionality per gene type.

### Exploring IGH Locus Allelic Diversity: Insights into Heterozygosity, and Novel Variants Across Individuals with Varied Ancestries

#### Allelic Distribution in the Studied Loci: Common and Unique Variants Among Individuals from Diverse Populations

To further explore allelic diversity at the IGH locus, we grouped the studied assemblies based on their broadly defined ancestral backgrounds, as African, Asian, European, Puerto Rican, Saudi, and mixed (African, European, Native American) (Table 1, Supplementary Table S1, sheet: Haploid Assemblies Support), and performed a Venn analysis (https://bioinformatics.psb.ugent.be/webtools/Venn/) (Figure 3). This approach revealed both widely shared alleles and others observed only in specific assemblies, reflecting the locus’s diversity within this dataset.

**Figure 3.**
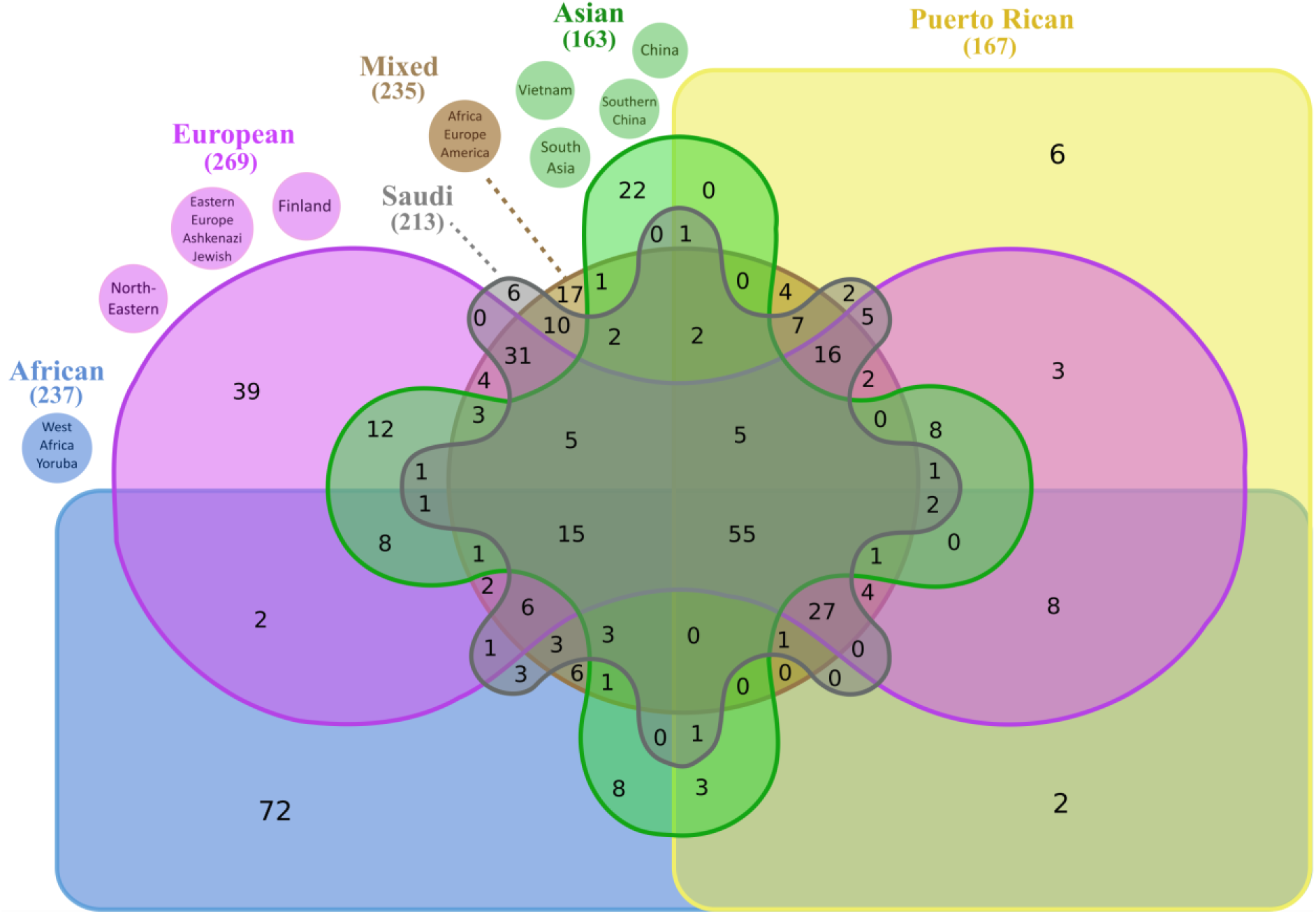
Genetic overlap and diversity: Venn diagram of IGH allelic distribution across assemblies of individuals from different ancestral backgrounds. This Venn diagram illustrates the distribution of alleles of the studied IGH loci in high-quality haploid assemblies from individuals of diverse populations. Each color represents assemblies associated with a given ethnicity background: brown for mixed (African, European, and Native American), green for Asian, gray for Saudi, yellow for Puerto Rican, pink for European and blue for African. The numbers in the intersections highlight alleles shared among assemblies from different population groups or identified exclusively in one, offering a visual insight into genetic overlap and divergence. The numbers in parentheses show the unique alleles identified across individuals of the same population. **Note:** This figure reflects only the individuals included in the current dataset and should not be interpreted as representative of a given population.

Fifty-five alleles, located within V-CLUSTER, were shared across all six groups, each found in at least one individual per group, underscoring a core set of conserved alleles. One additional allele, IGHV3-30-22*01 (located in CNV3) was present in five of the six groups (one African individual, one Asian, one European, one mixed, and one Puerto Rican assembly) and another 27 alleles, including several IGHC (IGHA2*01, IGHG4*01, IGHE*02) and one IGHD (IGHD1-1*01), were shared among 4 of the 6 groups, the studied African, European, mixed, Puerto Rican, and Saudi assemblies.

Examining the allelic relationships in more detail, several alleles were observed across loci of different ancestries. For example, IGH loci from the studied African individual shared 3 alleles with one Asian and one Puerto Rican individual, and 8 alleles with two assemblies of European and Puerto Rican ancestry. Eight IGHV alleles were shared among one African, two Asian, and four European individuals. In addition, one African and two Asian individuals shared 8 additional alleles, uniquely found in those three assemblies and two alleles were shared only between the African individual and one European assembly.

Some alleles appeared unique to individuals from a given background: 72 alleles were exclusive to the African individual, 39 to the European studied assemblies, 22 to the Asian assemblies, 6 to the Puerto Rican individual and 6 to the maternal and paternal Saudi assemblies (Supplementary Table S2). While the number of individuals per ancestry group is limited and these findings cannot currently be extrapolated to the population, our results confirm the high level of allelic polymorphism of IGH genes and highlight the important individual variation within the IGH locus.

A detailed breakdown of the functional IGHV allele frequencies across the studied assemblies (Table 1, Supplementary Table S1, sheet: Haploid Assemblies Support) is presented in Figure 4. Certain genes, such as IGHV3-43D and IGHV2-5, exhibited a relatively even distribution across the assemblies, while others showed a clear dominance of specific alleles. For instance, the IGHV1-18 gene displayed two alleles, with the *01 allele present in 90% of the studied assemblies, and the *04 allele observed only once, in a European sample. IGHV4-28 was predominantly represented by the *01 allele, which appeared in 69% of assemblies, followed by the *02 that was found twice, once in a Saudi individual and once in a European individual. The *05 and *07 alleles were each identified once, in an Asian and an African individual, respectively.

**Figure 4.**
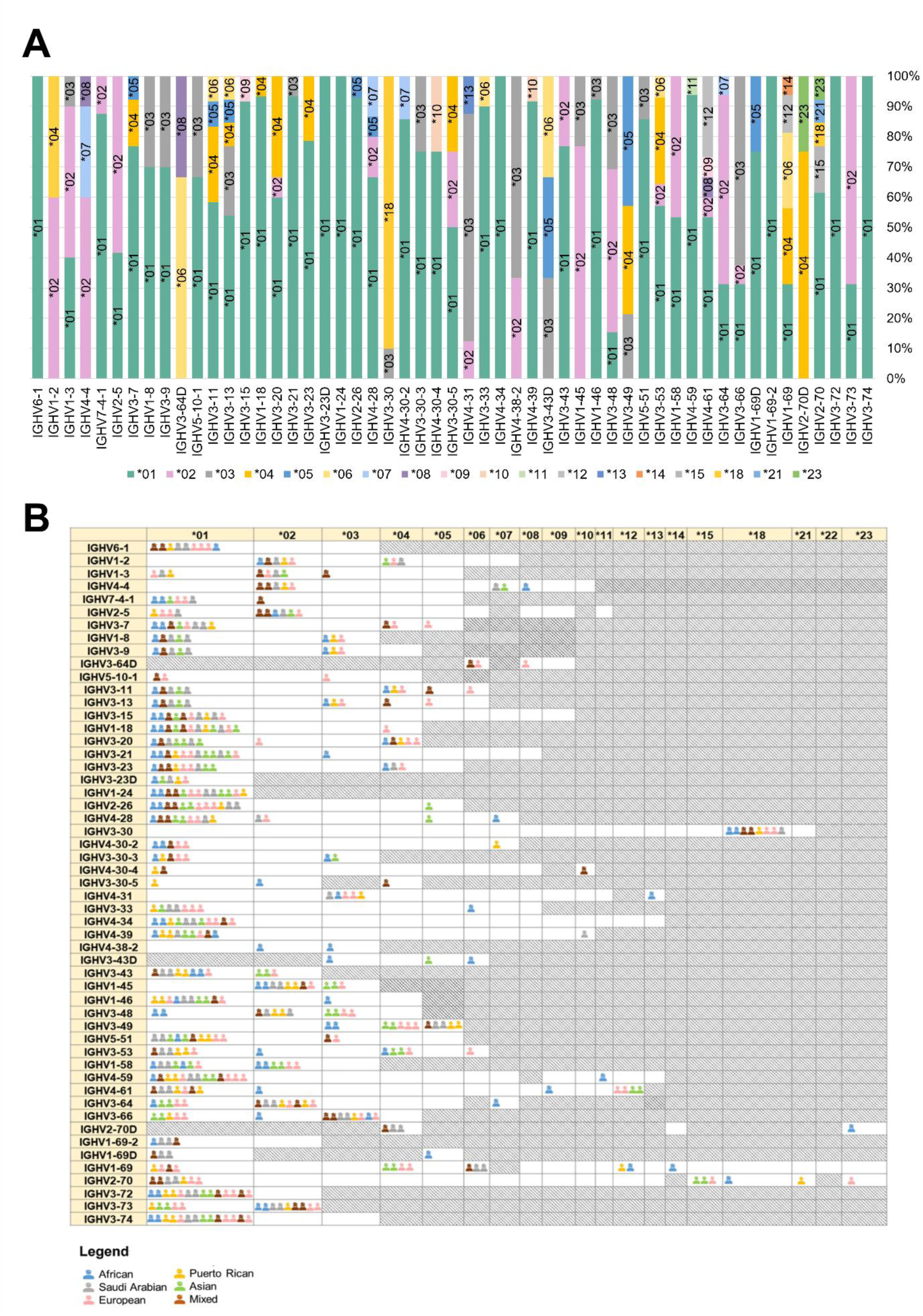
Functional IGHV Allele Frequencies and Gene-Specific Distributions. **(A)** The stacked bar chart illustrates the frequency distribution of functional IGHV alleles across the assemblies of our dataset. The allele frequencies (expressed as percentages) provide insights into IGHV gene variability and allele prevalence, highlighting patterns of diversity. Each allele is represented by a distinct color, with allele names labeled for clarity. **(B)** The table presents the distribution of IGHV alleles across the annotated assemblies with available population data. Grey cells indicate alleles that have not yet been found in a given IGHV gene, while white cells represent alleles that are present in the IMGT database but were not identified in the newly annotated assemblies that provide population information.

Among the analyzed assemblies, IGHV2-70 and IGHV1-69 appeared to be the most polymorphic genes, each exhibiting five distinct alleles. IGHV2-70 showed a distinct dominance of the *01 allele, which was present in 54% of the assemblies. The *15 allele was observed three times, specifically in the maternal and paternal Asian assemblies, as well as in an Eastern European individual. The remaining alleles—*18, *21, and *23—were each identified once in African, Puerto Rican, and European assemblies, respectively. For IGHV1-69, allele distribution was relatively even. The *01 and *04 alleles appeared four times each, the *06 allele three times, the *12 allele twice, and the *14 allele once. It should be noticed that IGHV1-69 and IGHV2-70 are among the known most polymorphic genes (globally 20 and 21 alleles respectively, in IMGT/GENE-DB, June 2025).

Notably, 10 IGHV genes (IGHV6-1, IGHV3-15, IGHV3-23D, IGHV1-24, IGHV2-26, IGHV3-30, IGHV4-34, IGHV1-69-2, IGHV3-72, and IGHV3-74) were present in only one allele, indicating both a lack of polymorphism and a high level of conservation across the assemblies analyzed.

### Heterozygosity at the IGH Locus in Relation to Individual’s Ancestral Diversity in Annotated Assemblies

A detailed exploration of heterozygosity within the IGH locus across individuals from diverse genetic backgrounds is presented in this study, evaluating allelic variation between their maternal and paternal genome assemblies. Heterozygosity, defined as the presence of two different alleles in genes shared between an individual’s maternal and paternal assemblies, was assessed by detecting only the common IGHV, IGHD, IGHJ and IGHC genes and quantifying the number of common and heterogeneous alleles, allowing us to gain insight into the degree of genetic diversity present at the IGH locus of each individual. It is important to note that several assemblies were derived from B lymphocytes, which undergo DJ or VDJ recombination. As a result, these assemblies may lack portions of the IGH locus. Therefore, heterozygosity percentages in these cases may be underestimated and are not directly comparable to those from non-rearranged samples. Nevertheless, calculating the proportion of heterozygous alleles within shared genes offers a meaningful view of IGH allelic diversity at the individual level, and complements prior studies that have emphasized the importance of heterozygosity in influencing immune response (43). Our results, illustrated in Figure 5, contribute to the understanding of IGH variation within individuals and mark an important step forward in characterizing immune gene diversity among individuals, while also emphasizing the need for broader investigations to confirm and expand upon our observations.

**Figure 5.**
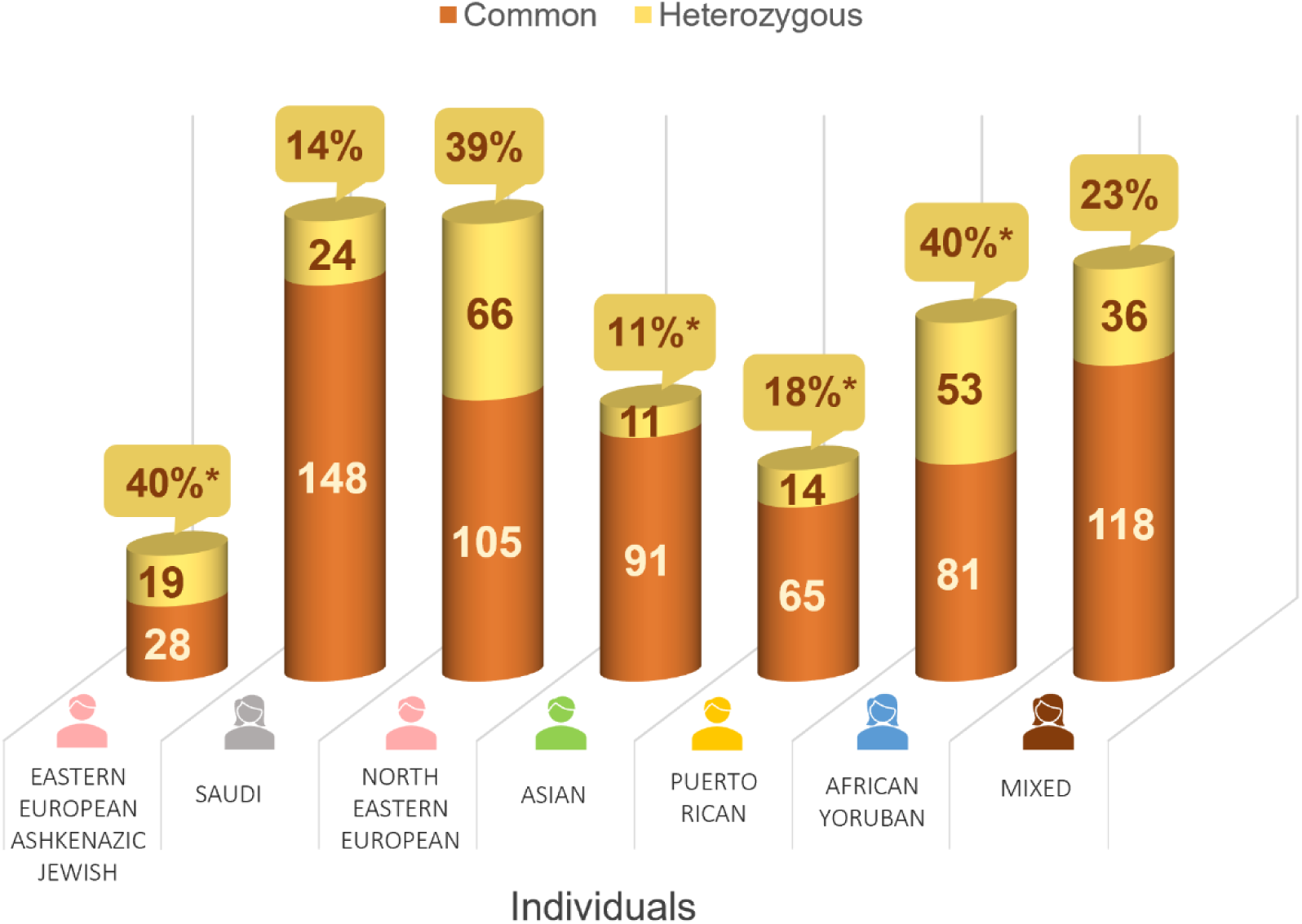
Heterozygosity analysis of the IGH locus across individuals with diverse ancestry. This graph illustrates the distribution of allelic variation within shared IGHV, IGHD, IGHJ, and IGHC genes in individuals for whom maternal and paternal assemblies are available. Only genes present in both assemblies were compared to assess allele concordance. Bars represent the total number of shared genes per individual: common alleles (identical in both assemblies) shown in orange, and heterozygous alleles (different between the two assemblies) shown in yellow. The percentage of heterozygous alleles per individual is indicated above each bar; asterisks denote cases where one or both assemblies were derived from rearranged B cells and may lack intact IGH genes due to DJ or VDJ recombination (more details available at: https://www.imgt.org/IMGTrepertoire/index.php?section=LocusGenes&repertoire=GeneOrder&species=human&group=IGH). In such cases, heterozygosity should be interpreted with caution. Population-specific colors are consistently used across all figures in this study, and the population description is respected based on the sample information. **Note:** This graph presents heterozygosity data from seven individuals and is not intended to represent population-level heterozygosity.

### Novel Alleles and their Distribution among Individuals of Varied Genetic Backgrounds

A total of 120 novel IGH alleles were identified across the analyzed assemblies, including 84 (14 F, 5 ORF and 65 P) IGHV alleles (53 belonging to subgroups and 31 classified in clans), 3 IGHD (2 F and 1 ORF), 2 IGHJ (1 F and 1 P) and 31 IGHC alleles (22 F, 4 ORF and 5 P), including the novel identification of the IGHG4A gene (Supplementary Table S3 and Supplementary Figure S3). The highest number of novel alleles was detected in assemblies of the African individuals (36 alleles), followed by those of European (30 alleles) and Asian individuals (22 alleles). Notably, the reference assembly BK063800 (T2T-CHM13v2.0) contributed the 3 newly identified IGHD alleles, while the IGHJ set was enriched with two alleles, one allele (IGHJ5*03 P) found in the African Yoruba assembly (NA19240.pri.mat.f1_v2) and one allele (IGHJ5*04 F) in the Caucasian assembly (hg002v1.0.1.mat). Several newly described alleles were shared among the assemblies of various genetic background, with 4 IGHV alleles shared between the Asian and European individuals, and 3 IGHC alleles found across European, mixed, Saudi and Puerto Rican (Figure 6, Supplementary Table S4).

**Figure 6.**
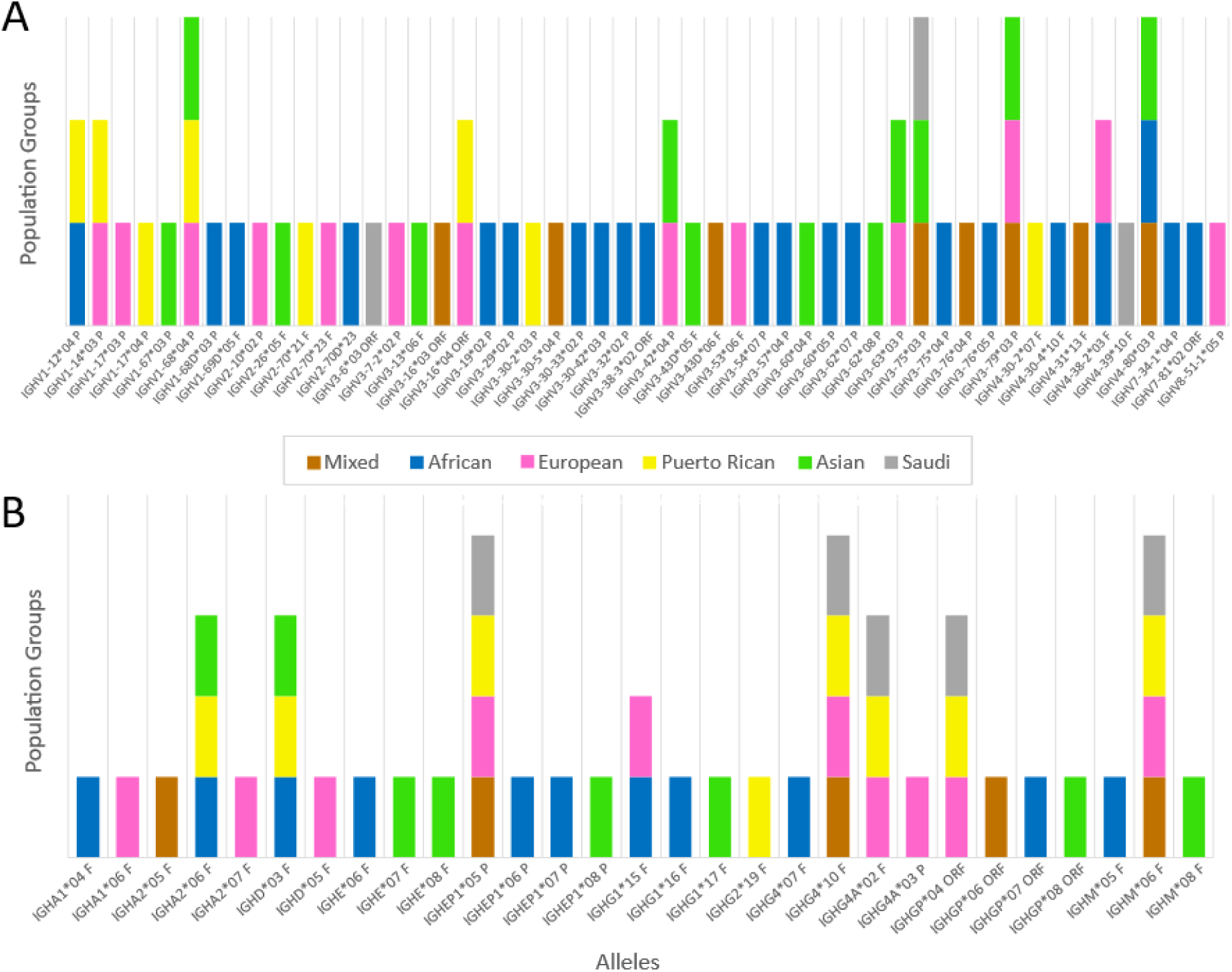
Distribution of novel alleles at the IGH locus among annotated assemblies from different population groups. This bar graph illustrates the distribution of novel IGHV and IGHC alleles identified across assemblies of African, European, Asian, Puerto Rican, Saudi, and mixed ancestries. Each colored bar represents a different population group. To avoid redundancy, each unique allele is counted only once for each ancestry group, regardless of the number of individuals it appears in. **(A)** Novel IGHV alleles: The novel allele IGHV3-15*09 (IMGT000112, JADEBP010005792) is not included due to the absence of information regarding the samples’ ethnicities. Alleles identified in complementary scaffolds (e.g. JADDCI010010883, JADCYG010001029, QEHY01024525, JADDFX010000192) are included. Note: This graph shows only IGHV alleles that belong to subgroups, as per the classification used in the study. For clans and further information, see Supplementary Table S4. **(B)** Novel IGHC alleles: The novel alleles IGHM*07 (BK063802), IGHG4*09, and IGHGP*04 (BK063800) are not included in the graph, due to unavailable population information or lack of assembly correspondence to a specific individual.

### IMGT Database Enrichment: Updates and Functional Enhancements to the IGH Locus Dataset

#### IMGT Human IGH Reference Dataset: Enrichment with Novel Alleles for Enhanced Diversity

The IMGT/GENE-DB reference dataset for the IGH locus (IMGT/GENE-DB program version: 3.1.42, 2024-09-25) has been enriched significantly, now comprising a total of 264 genes and 872 alleles (Table 2). This update notably enhances the dataset for the IGH locus, surpassing the IMGT reference dataset used in the beginning of the study (version 3.1.38, 2022-11-08) by incorporating in total 124 novel alleles. Of these, 120 were identified in newly annotated assemblies and supporting scaffold annotations, and 4 IGHV genes were identified in existing IMGT sequences. This enrichment is characterized by the addition of 88 novel IGHV alleles (17% increase), along with 3 new IGHD alleles, and 2 new IGHJ alleles. The most notable advancement is in the IGHC set, which has been expanded by 34%, incorporating 31 new alleles, including the remarkable addition of a newly identified and annotated IGHC gene, the IGHG4A. Table 2 provides a summary of the allele numbers and percentage increases of the updated IMGT IGH reference dataset, including genes located on chromosome 14, as well as non-localized genes (genes identified but not assigned to a specific chromosomal position) and orphons (genes outside the main locus) (https://www.imgt.org/IMGTindex/orphon.php/). Further details on the newly added alleles are available in Supplementary Table S3.

**Table 2.**
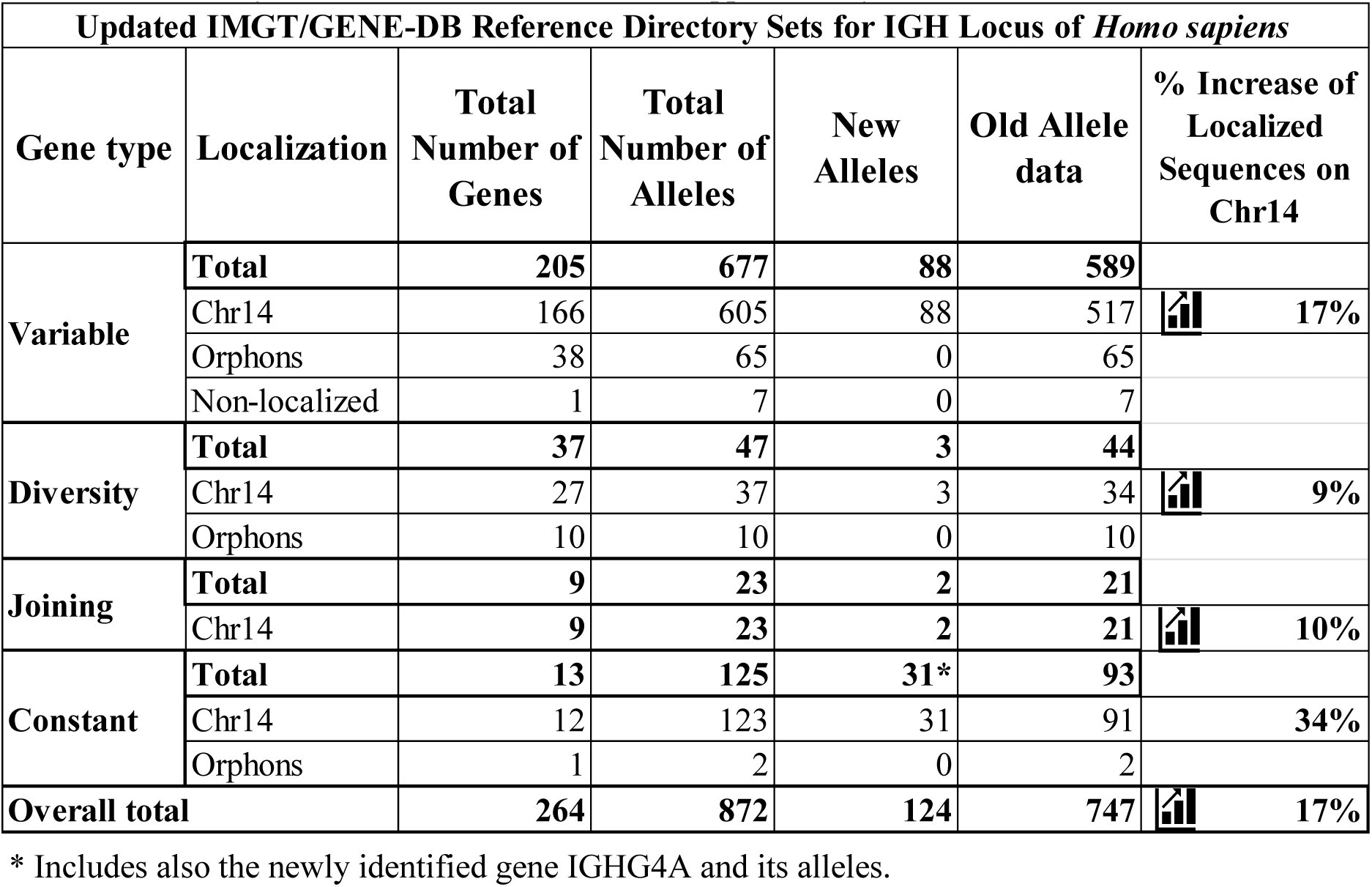
Updated IMGT/GENE-DB Reference Directory Sets for IGH locus of *Homo sapiens*.

|  |  |  |  |  |  |  |
| --- | --- | --- | --- | --- | --- | --- |
| Gene type | Localization | Total Number of Genes | Total Number of Alleles | New Alleles | Old Allele data | % Increase of Localized Sequences on Chr14 |
| Variable | Total | 205 | 677 | 88 | 589 |  |
|  | Chr14 | 166 | 605 | 88 | 517 | 17% |
|  | Orphans | 38 | 65 | 0 | 65 |  |
|  | Non-localized | 1 | 7 | 0 | 7 |  |
| Diversity | Total | 37 | 47 | 3 | 44 |  |
|  | Chr14 | 27 | 37 | 3 | 34 | 9% |
|  | Orphans | 10 | 10 | 0 | 10 |  |
| Joining | Total | 9 | 23 | 2 | 21 |  |
|  | Chr14 | 9 | 23 | 2 | 21 | 10% |
| Constant | Total | 13 | 125 | 31* | 93 |  |
|  | Chr14 | 12 | 123 | 31 | 91 | 34% |
|  | Orphans | 1 | 2 | 0 | 2 |  |
| Overall total |  | 264 | 872 | 124 | 747 | 17% |

\* Includes also the newly identified gene IGHG4A and its alleles.

#### IMGT Updates: Replacement of Partial Sequences

The updated IMGT/GENE-DB reference dataset (IMGT/GENE-DB program version: 3.1.42, 2024-09-25) is now accessible through dynamic gene tables (https://www.imgt.org/IMGTrepertoire/LocusGenes/genetable/autotable.php, version: v.1.30) that provide comprehensive information about each gene and allele (including whether they are fully characterized or partial in the reference data). In these updated tables, sequences missing a complete X-GENE-UNIT (X stands for L-V, D, J or C) are indicated by an asterisk.

Among the 599 IGHV gene alleles located on chromosome 14 (excluding the 108 highly truncated alleles which belong to clans), 188 (38.2%) are flagged as incomplete. Notably, only three of these, IGHV1-12*04 P (IMGT000176), IGHV1-14*03 P (IMGT000176) and IGHV4-80*03 P (IMGT000111), are newly incorporated, featuring a complete V-REGION but lacking the V-RS. Of the 188 incomplete sequences, without a complete L-V-GENE-UNIT, 76 (∼40.4%) have partial V-REGION due to incomplete sequence availability (partial in 5’ or 3’ or both). In contrast, the remaining 112 (∼59.6%) possess complete V-REGION, but are missing critical components, such as leader sequences (L-PART1, L-PART2) and/or recombination signal sequences (V-HEPTAMER, V-NONAMER).

As part of our analysis, we replaced in total 43 partial sequences with their corresponding complete ones. More specifically, 24 IGHV partial allele sequences, previously lacking L-V-GENE-UNIT and/or exhibiting incomplete partial V-REGION, were replaced by complete sequences identified in the new assemblies. Furthermore, to enhance the accuracy of our data and results, we revised the annotations of accession numbers that were already present in the IMGT databases. This revision led to the replacement of 7 partial sequences with complete sequences, as well as the addition of 4 novel alleles, all associated with existing accession numbers.

Regarding the IGHC genes, this analysis replaced 15 partial constant exon sequences with complete ones, of which 13 were identified through this new assembly analysis and 2 were corrected using existing complete sequences from the IMGT databases. This update resulted in a 20% increase in the number of complete constant gene sequences within the dataset, and remarkably, all new constant alleles introduced in this study are complete, with the exception of the IGHEP1 alleles, which are inherently lacking CH1 and CH2 exons.

Detailed information on sequence completeness and updates is provided in Supplementary Table S3 and the corresponding dynamic gene tables.

### Updated IMGT Reference Set: New Reliability System for IGH Alleles

The confidence in the reliability of the alleles within the updated IMGT dataset is quantitatively assessed through the IMGT allele confirmation score, which utilizes a star rating system ranging from 1 to 3 stars. This scoring system is available in IMGT’s dynamic gene tables (https://www.imgt.org/IMGTrepertoire/LocusGenes/genetable/autotable.php) and reflects increasing confidence levels in the existence of each allele, based on the available supporting evidence; literature sequences previously integrated in IMGT and relevant publications. In addition, the novel alleles identified in this study have been confirmed using the methodology described in section *Evaluation of Newly Identified Alleles for Accuracy and Reliability* and are listed in Supplementary Table S5.

Figure 7 illustrates the distribution of the updated IGH human IMGT/V-QUEST reference dataset (202518-3, 30 April 2025), per functionality and confidence level. Alleles classified as ’high confidence’ are those supported by at least one literature sequence, earning an IMGT allele confirmation score of 2 or 3 stars, or newly identified alleles lacking literature sequence support but validated by methodology. The remaining alleles are classified as ’low confidence’.

**Figure 7.**
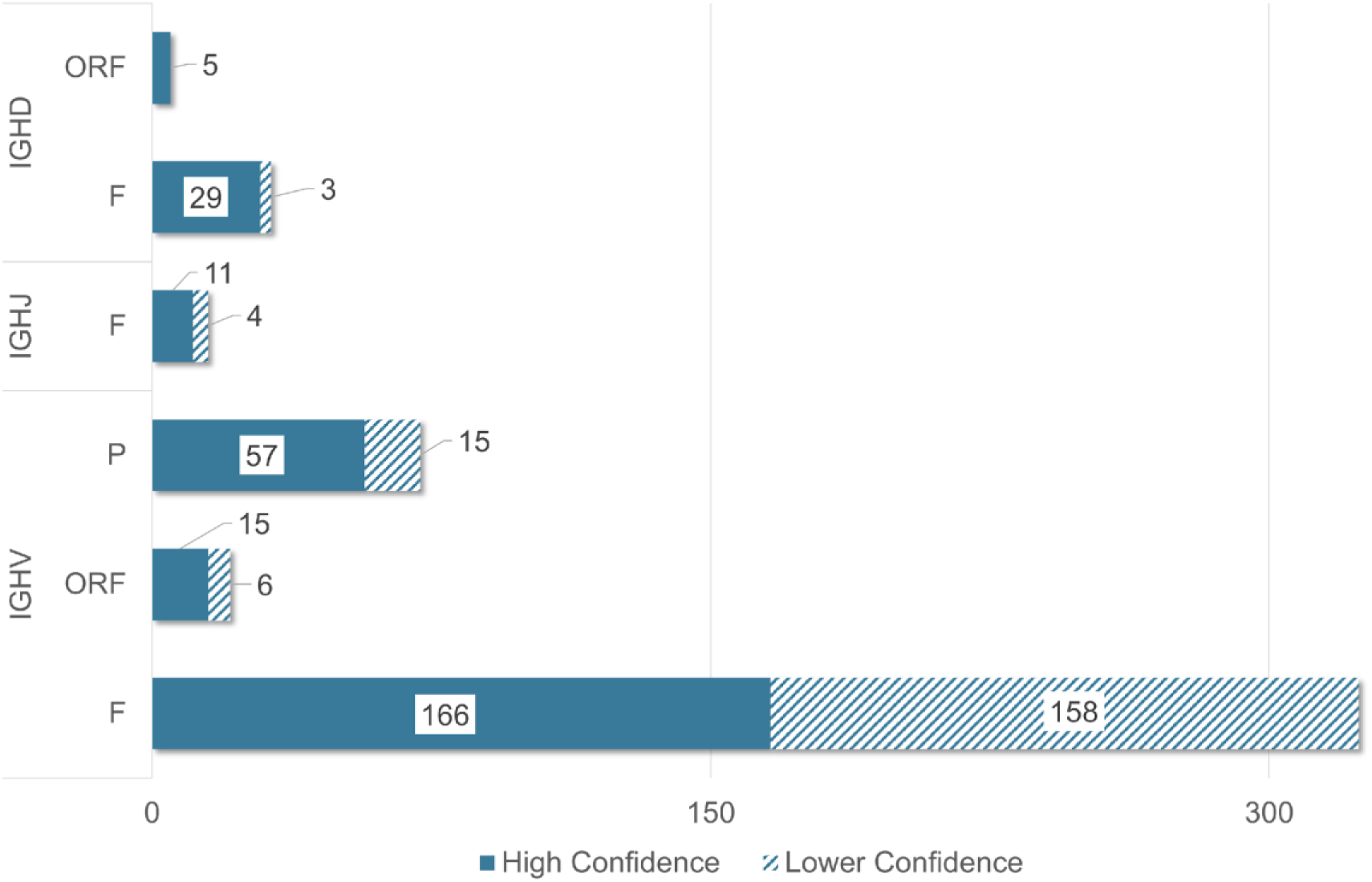
Distribution of IMGT/V-QUEST reference dataset (202518-3, 30 April 2025) by gene type, functionality, and confidence level. The chart illustrates counts of F, ORF, and P for IGHV, IGHD and IGHJ alleles. Confidence levels are classified as ’High’, for alleles with more than 2 IMGT allele confidence scores or newly identified 1-star alleles validated by methods of 2.5, and ’Lower’ (for 1-star alleles). Orphons are excluded.

The updated IMGT reference set includes 417 IGHV alleles (327 F, 21 ORF, and 72 P), with 57% classified as ’high confidence’. Conversely, 179 IGHV alleles in the dataset have received a one-star IMGT allele confirmation score; among these, 41 are also present in the OGRDB reference set (44), thereby increasing their confidence level.

As for the constant region, the updated IMGT/GENE-DB reference set includes 12 IGHC genes, which collectively comprise 121 different alleles (Figure 8A). Of these alleles, 104 are classified as F, 6 as ORF, and 11 as P (Figure 8B). Notably, 56% of the sequences have been classified as ‘high confidence’ as they have been identified multiple times within the dataset, resulting in confidence scores of 2 or 3 asterisks or have been confirmed using the previously mentioned validation methods, as detailed in Figure 8A.

**Figure 8.**
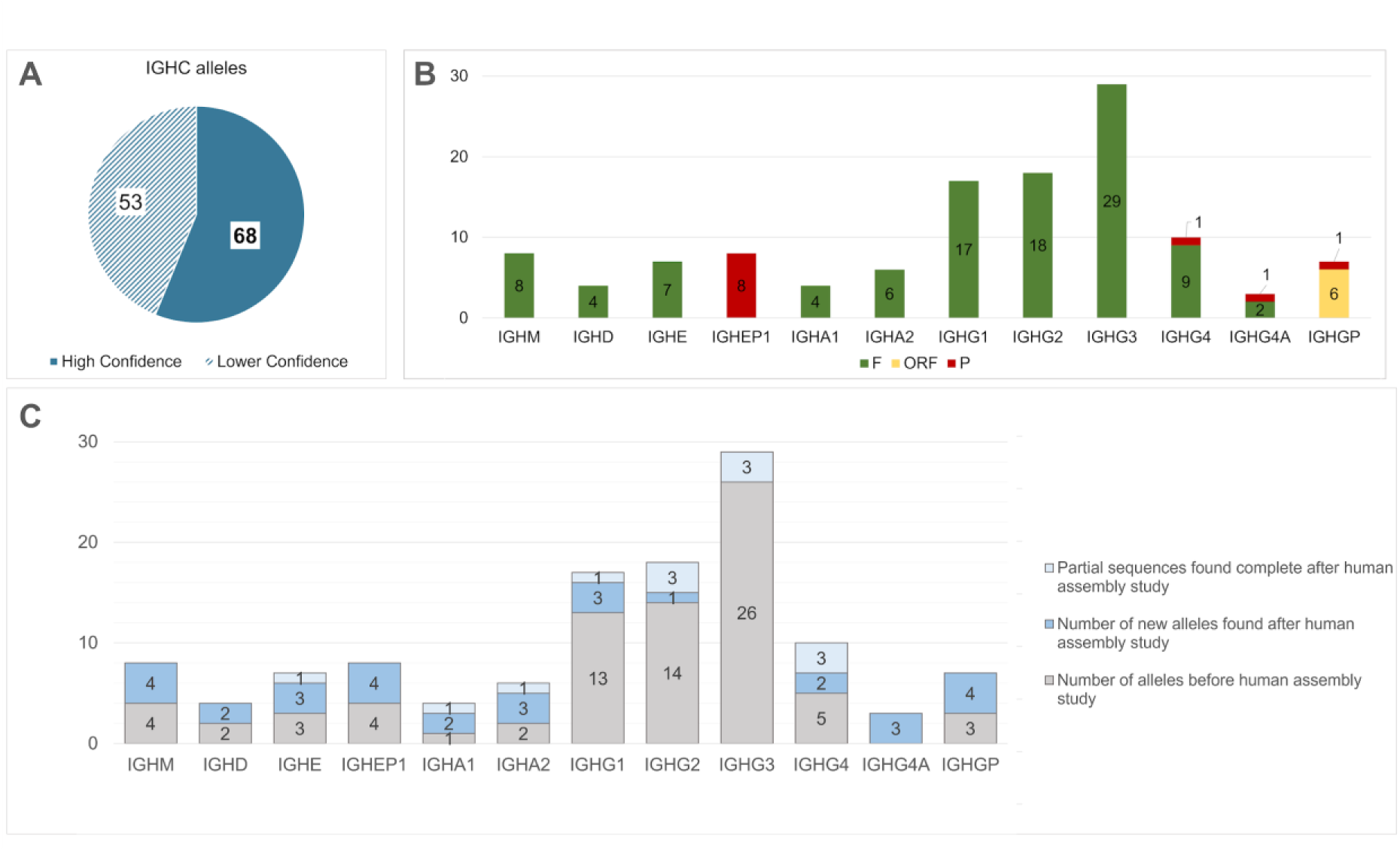
Updated IMGT/GENE-DB IGHC reference set. **(A)** Confidence level breakdown for IGHC. Confidence levels are classified as ’High’, for alleles with more than 2 IMGT allele confirmation scores or newly identified 1-star alleles validated by methods of 2.5, and ’Lower’ (for 1-star alleles). **(B)** Distribution of IGHC classes and subclasses of IMGT reference set per functionality. **(C)** Allele number distribution before and after the human assembly study, showing the number of partial genes found complete, newly identified genes and previously known genes, for each class/subclass.

The current analysis enriched the IMGT dataset by adding new alleles and completing previously partial alleles where certain exons were missing (Figure 8C). A total of 31 novel alleles were detected, and 13 partial alleles were found to be complete (Supplementary Table S3). This results in a 55.7% overall enrichment of the dataset. As shown in Figure 8C, the highest number of new alleles were found in IGHGP, with 4 out of 7 alleles (57%) being newly identified. IGHM, IGHD, IGHE, IGHA1, IGHA2, and IGHEP1 followed, each exhibiting 50% novel alleles.

### IMGT CNV and haplotype Diversity in the IGH Locus: Definitions, Observed Patterns, and Unique Rearrangement Across Individuals from Diverse Backgrounds

#### Defining IMGT CNV Forms and IMGT haplotypes

Copy Number Variations (CNVs) refer to structural alterations in the genome, such as deletions, insertions, or duplications, that change the gene copy number. Each IMGT CNV is characterized by a functional gene at its 5’ end and another at its 3’ end (7). The variations in gene copy number within a CNV are classified as IMGT CNV forms, each denoted by an alphabetical letter, starting from ’A’. The unique combinations of IMGT CNV forms in an individual contribute to the characterization of their IMGT haplotype for the IGH locus (Figure 9).

**Figure 9.**
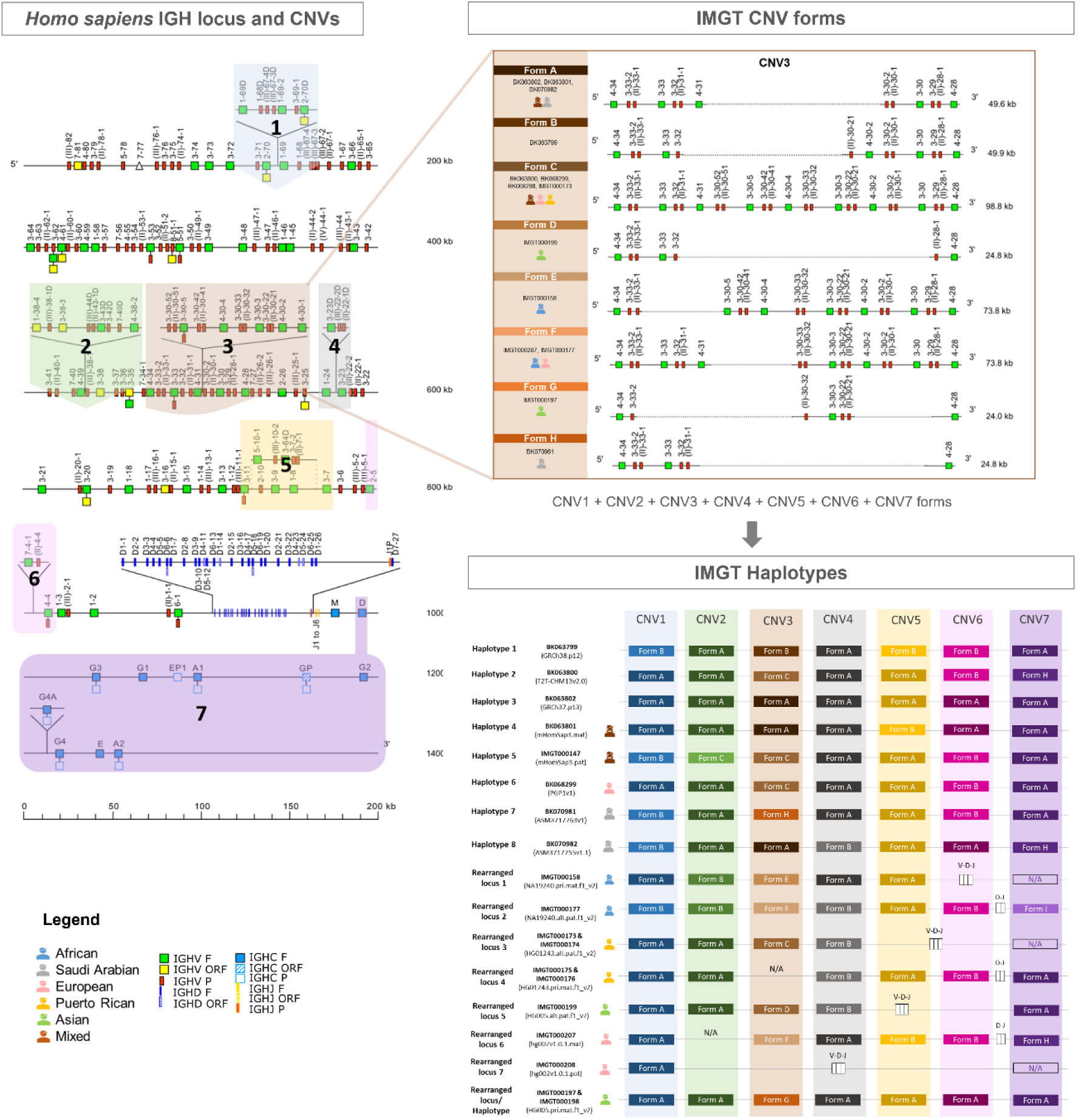
Schematic illustration of the IMGT concepts of CNVs, IMGT CNV forms and IMGT haplotype. Each CNV can have multiple variations, referred to as CNV forms. The Figure illustrates all 7 CNVs in the human IGH locus and the way we analyze the different forms of each CNV. The combination of these forms across different CNVs in a given locus constitutes an individual’s IMGT haplotype.

### IMGT CNV forms Analysis: Initial Insights into Characteristics and Patterns Across Individuals of Diverse Population Groups

Within the human IGH locus, seven CNVs have been identified: six involving IGHV genes and one involving IGHC genes.

CNV1, CNV2, and CNV4 each displayed variants characterized by gene duplications and insertions (Figure 10). Form A, which represents the non-duplicated version of the genes of these CNVs, appeared to be the most prevalent across all studied genome assemblies. The duplicated form, form B, of CNV1 was observed in an African and a mixed African-European-Native American sequence, while form B of CNV4 was noted in four assemblies of Asian, Puerto Rican, African, and Saudi individuals. CNV2’s duplicated form B was only detected in the maternal and paternal IMGT haplotypes of the African assembly, while form C, lacking the first four genes, occurred only once in the mixed assembly.

**Figure 10.**
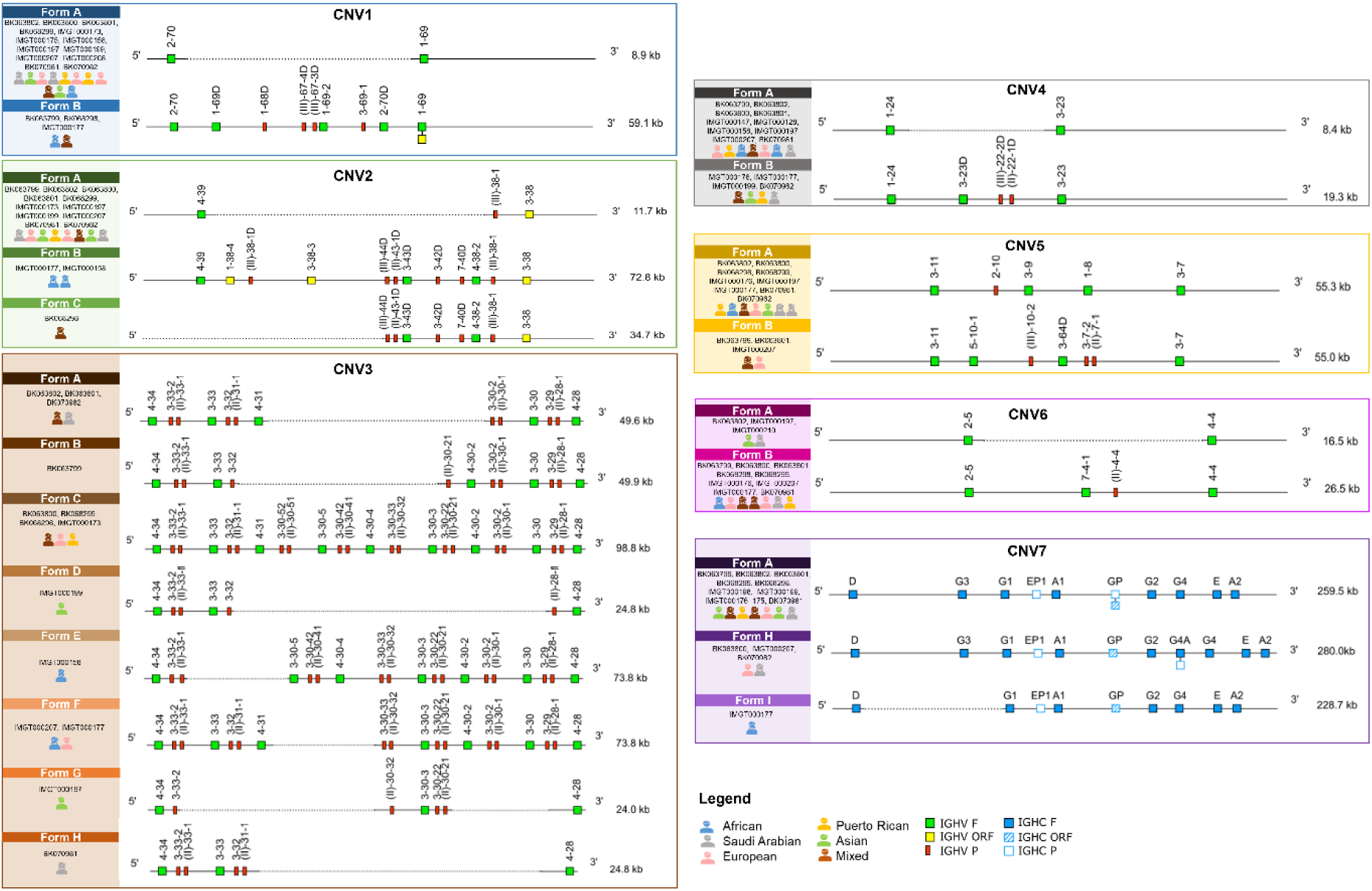
Comprehensive analysis of IGH IMGT CNV forms in human individuals across different ethnic groups. Detailed overview of the 7 human IGH CNVs and their IMGT CNV forms, including the corresponding sequences and the ethnicities in which each form was identified. All findings of the CNVs and their forms derived from the current analysis can be accessed on the updated IMGT website under ‘Copy Number Variations (CNV) forms in human *(Homo sapiens)* IGH’ (https://www.imgt.org/IMGTrepertoire/index.php?section=LocusGenes&repertoire=locus&species=human&group=IGH).

CNV3 stood out as the most polymorphic region, displaying high variability due to multiple duplications of genes of IGHV3 and IGHV4 subgroups, posing challenges for description and annotation. The current study revealed eight forms of CNV3, each of which was observed between one and four times, without any of them showing high dominance. Form C, which was detected most frequently (four times), includes all known genes associated with CNV3.

Moreover, CNV5 consists of two forms that contain the same number of functional genes, though gene classification and their distances differ. Form A is found in most sequences, while form B is observed three times: in the reference sequence GRCh38.p14, in a mixed and a European assembly.

In parallel, CNV6 also has two forms. Form B includes two additional genes (one F and one P) compared to form A, with form B identified in nine sequences and form A in three.

Finally, CNV7 pertains to the IGHC genes with three distinct IMGT CNV forms identified. The dominant form, A, includes all IGHC genes without the IGHG4A duplication and is observed in nine sequences. Form H, which is characterized by the IGHG4 duplication, is present in three sequences, while the form I is only observed once.

### Identification of Distinct IMGT haplotypes and Rearranged Loci

The current research described 8 distinct IMGT haplotypes of the IGH locus across 8 assemblies, resulting from the combination of various CNV forms (Figure 11). Sequences with detected rearrangements, where some of IGHV, IGHD, or IGHJ genes were missing, were classified as IMGT Rearranged Loci instead of haplotypes. Eight rearranged loci with unique CNV form combinations were annotated, providing insights into potential new IMGT haplotypes. Notably, no combination of IMGT CNV form was found more than once.

**Figure 11.**
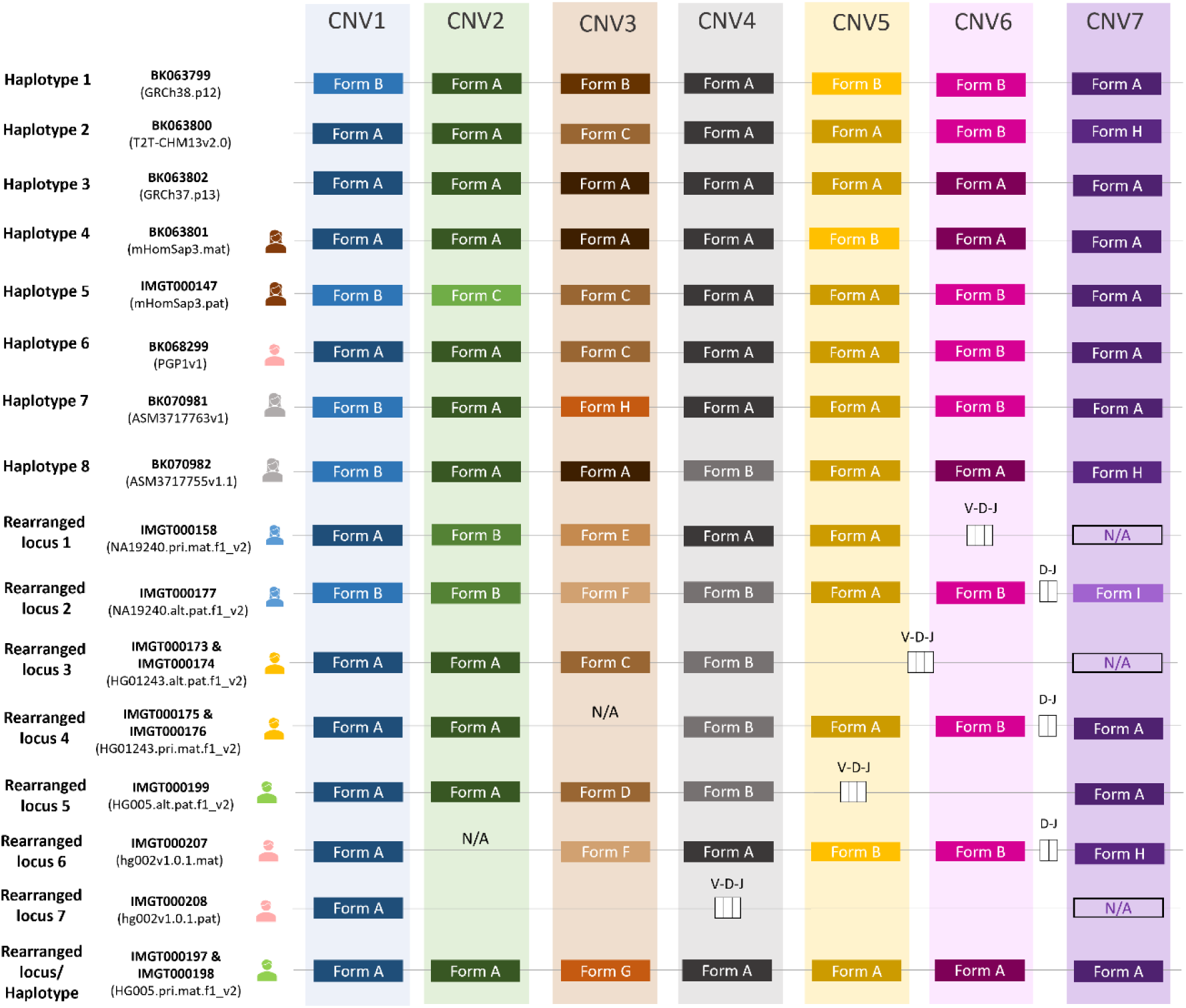
Comprehensive overview of IMGT haplotypes and rearranged loci details across annotated human genome assemblies. This figure provides a detailed analysis of IMGT human haplotypes and rearranged loci for each annotated assembly. N/A stands for the CNV forms that could not be annotated; CNV2 for rearranged locus 6 remains unidentified due to missing genes between IGHV4-59 and IGHV(III)-38-1, CNV3 for rearranged locus 4 is not annotated due to discrepancies between the two corresponding scaffolds and three CNV7 cases due to potential class switch. White boxes indicate rearrangements within the loci as specified by the IMGT standard coloring menu (https://www.imgt.org/IMGTScientificChart/RepresentationRules/colormenu.php).

### Comprehensive Analysis of IGHC genes

#### Identification of IGHG4 duplication: Gene IGHG4A

A significant result of the current study is the identification and annotation of a novel IGHC gene, IGHG4A, which is a duplication of IGHG4. This duplicated gene is located approximately 15.2 kb downstream of IGHG2 and 15.4 kb upstream of IGHG4 (Figure 12). Notably, the distances between these genes do not vary significantly; in assemblies without the duplication, the average distance between IGHG2 and IGHG4 is 14.9 kb, while for IGHG4 to IGHE in non-duplicated cases, the distance ranges from 20.1 to 22.7 kb, with median of 20.1 kb, which aligns with the distance observed in the duplicated IGHG4 (Figure 12A).

**Figure 12.**
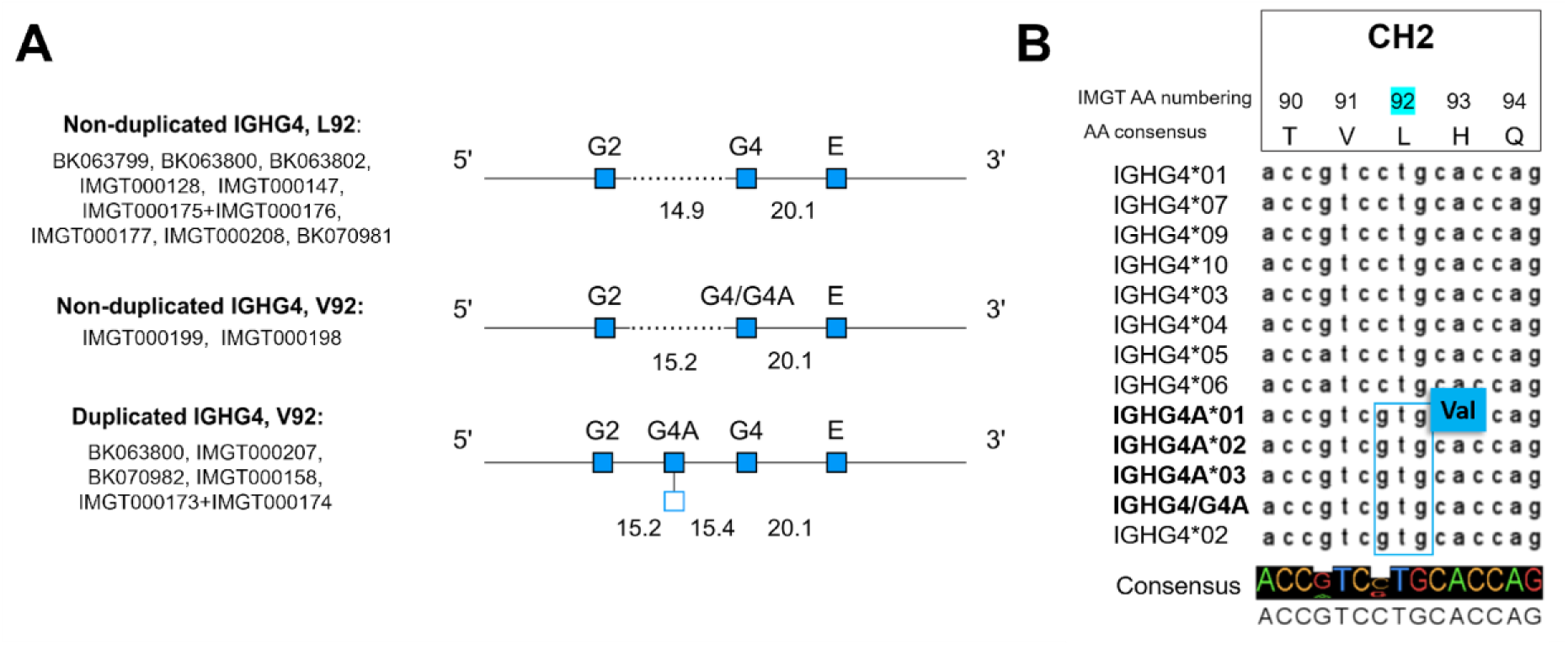
Gene structure and polymorphism analysis of IGHG4: duplication, allelic variation, and amino acid substitution. **(A)** Gene structure representation of IGHG4 duplication in assemblies: representation of non-duplicated and duplicated sequences with L92 and V92 variants. **(B)** MSA analysis of IGHG4 and IGHG4A alleles. Polymorphisms responsible for the V92 amino acid substitution are highlighted. MSA was calculated with MAFFT (31) and visualized in Jalview (46). Alleles sequences were derived from IMGT/GENE-DB.

This duplication was observed in five of the 15 analyzed assemblies (31.25%), represented by three alleles: two classified as F and one classified as P due to a frameshift in the hinge region.

IGHG4A is characterized by an amino acid substitution at position 92 within the CH2 exon, where leucine (L) is replaced by valine (V), confirming previous studies (45) (Figure 12B). In the current analysis, a non-duplicated IGHG4 gene with the V92 substitution was identified in two assemblies from an Asian individual (IMGT000199 and IMGT000198) (Figure 12B). In total, 11 sequences with non-duplicated IGHG4 were identified. Of these, 9 sequences (56.25%) carry the L92 variant, while 2 sequences (12.5%) harbor the V92 mutation, designated as IGHG4/G4A, due to the close similarity with the IGHG4A. Interestingly, both V92 mutations were found in the Asian individual, who was homozygous for the mutation (present in both maternal and paternal assemblies).

### Allele Frequency Distribution and Variability Patterns in IGHC genes

Figure 13 highlights the variability of each subclass and their allele frequencies in IMGT/GENE-DB. IGHG subclasses show the highest polymorphism, with IGHG2 (18 alleles) dominated by IGHG2*02 (27%), with the other alleles ranging from 3% to 11%, while IGHG1 (17 alleles) is dominated by IGHG1*03 (34%). IGHG3 (29 alleles) has a more balanced distribution, with IGHG3*11 being the most common (16%), followed by IGHG3*03 (11%) and IGHG3*14 (10%), while the remaining alleles range between 1% and 7%.

**Figure 13.**
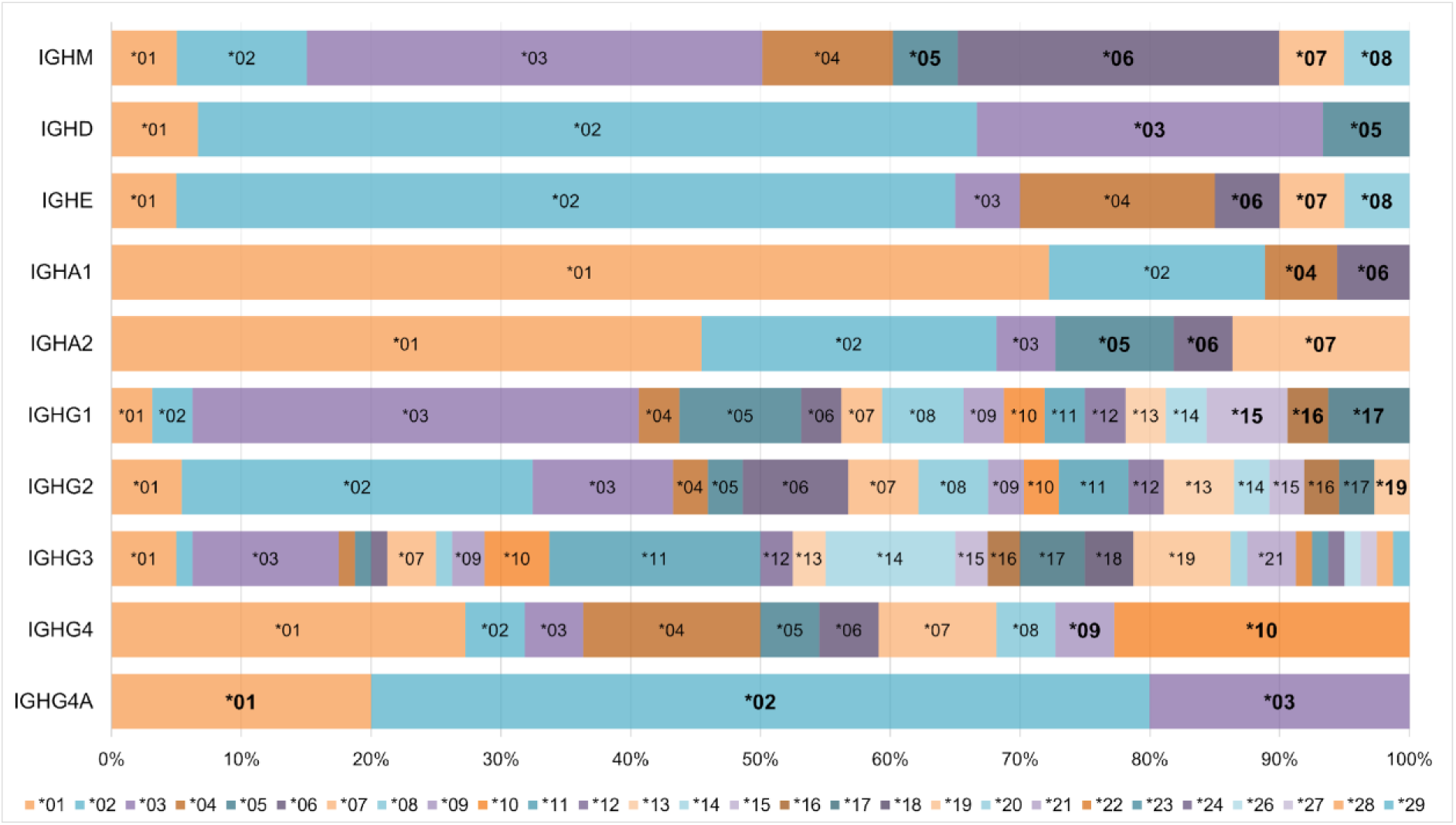
Stacked bar chart depicting the distribution of allele frequencies (in percentages) across each class and subclass of IGHC genes, providing a clear visualization of allele prevalence and distribution across different gene subclasses. Each allele is represented by a distinct color, with allele names labeled for those with frequencies greater than 1%. New alleles are highlighted in bold.

IGHE is dominated by IGHE*02 (60%), followed by the IGHE*04 (15%) and the rest with 5%. Regarding IGHA1, it is dominated by IGHA1*01 (72%), while IGHA2 is led by IGHA2*01 at 45%. Finally, IGHG4 (10 alleles), IGHG4*01 is the most common (27%), followed by the new IGHG4*10 (23%), indicating its potential significance within the population. Similarly, novel alleles in other gene classes also show relatively high frequencies. For instance, in the IGHM class, IGHM*03 is the most dominant allele, representing 37% of the sequences, followed by the novel IGHM*06 at 26%. In IGHD, the most prevalent allele is IGHD*02, accounting for 60%, with the novel IGHD*03 coming second with 27%. The high frequency of these novel alleles highlights the importance of these variants.

### Identification of Allotypes, new Polymorphisms and Variants in IGHC genes

This study uncovered novel polymorphisms leading to amino acid substitutions in IGHC genes where no allotypes have been described (IGHM, IGHD, IGHE, IGHA1 and IGHG4), and expanded the understanding of Gm and A2m allotypes through the identification of new alleles.

### IGHM, IGHD and IGHE

Three newly identified IGHM alleles display distinct variations that have not been previously documented, as shown in Figure 14A. IGHM*05, features a replacement of leucine to phenylalanine at position 21 (L21F) in the CH2 exon’s B-STRAND, observed in the African sequence IMGT000177; IGHM*07 (BK063802) shows a valine-to-glycine substitution at position 126 (V126G) in CH2; and IGHM*08 exhibits a threonine-to-isoleucine substitution at position 1.1 (T1.1I) in CH3, identified in the Asian sequence IMGT000198 (Figure 14A).

**Figure 14.**
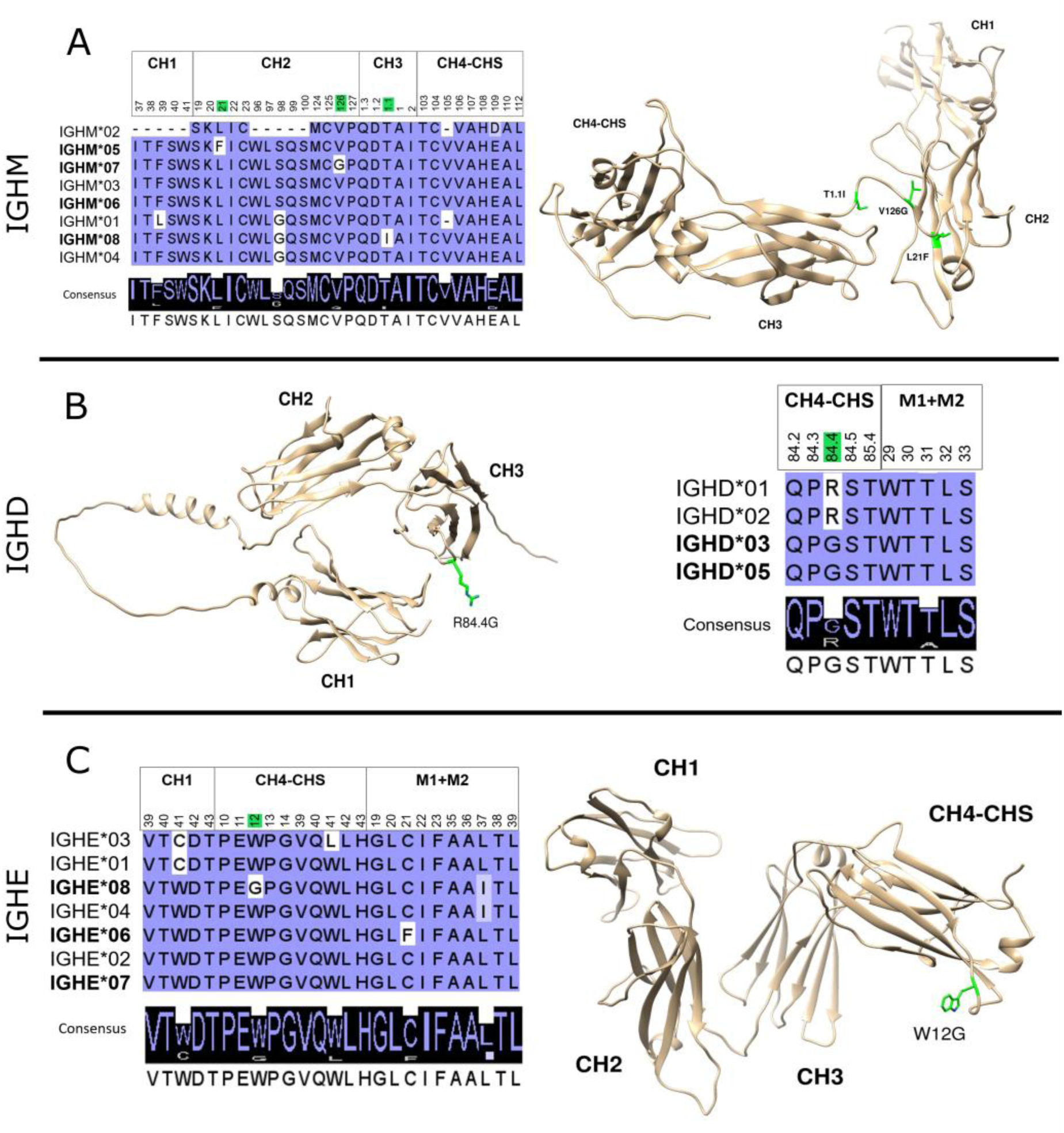
Novel mutations in the IGHM **(A)**, IGHD **(B)**, and IGHE **(C)** alleles, presented through MSA and structural protein representations. The models illustrate the constant region domains of IgM, IgD, and IgE. For IgM and IgE, the models display CH1, CH2, CH3, and CH4 domains, with CHS, the hydrophilic C-terminal end of secreted IG, located in 3’ of the CH4, while for the IgD features the CH1, CH2 and CH3 domains (47). The amino acid coding sequences were obtained from IMGT/GENE-DB, aligned using MUSCLE (v3.8.425), and visualized in Jalview with BLOSUM62 shading. The MSA figures display the regions with amino acid substitutions, with novel alleles in bold and the positions of the mutations marked in green. These mutations are also mapped green onto structural models of IgM (AF_AFP01871F1), IgD (AF_AFP01880F1), and IgE (AF_AFP01854F1), predicted by AlphaFold and visualized in Chimera (48).

The novel alleles IGHD*03, and IGHD*05, introduce a new substitution of G at position 84.4 in the CH3, replacing the arginine (R) (R84.4G) found in previous alleles (Figure 14B). This variation does not appear to be associated with any specific ethnicity as it is identified in individuals from diverse regions, including North-Eastern Europe (BK068299), Africa (IMGT000177), Asia (IMGT000198), and Puerto Rico (IMGT000176). Among the three new IGHE alleles (IGHE*06, IGHE*07, and IGHE*08), IGHE07 shares an identical amino acid sequence to IGHE*02, with silent mutations in the CH1 and CH2 exons. IGHE*06 of the African sequence IMGT000177, shows a novel cysteine-to-phenylalanine substitution at position 31 (C31F) in the transmembrane region. Meanwhile, IGHE*08, identified in an Asian assembly (IMGT000199), is distinguished by a G at position 12 in the CH4-CHS, replacing tryptophan (W), and an isoleucine at position 37 of the transmembrane region, also seen in IGHE*04 (Figure 14C).

### IGHA1 and IGHA2

IGHA1 lacks identified allotypes; however, a new variation has been identified in the allele IGHA1*06, of a North-Eastern European individual (BK068299). This variant features a mutation at position 95 of CH2, where glutamic acid (E) is replaced by aspartic acid (D) (E95D), indicating a rare variation within IGHA1 Figure 15.

**Figure 15.**
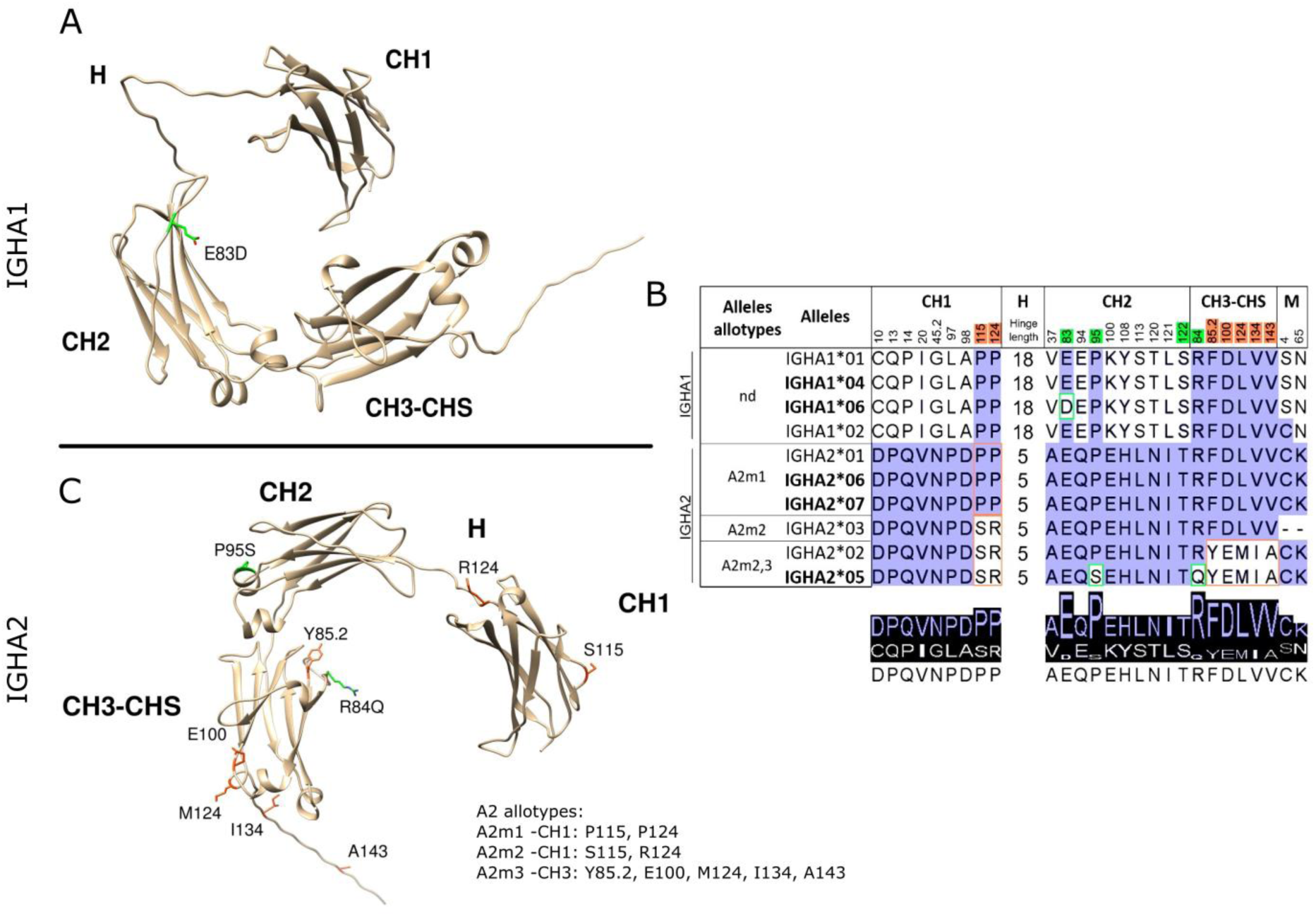
Novel mutations in the IGHA1 and IGHA2. **(A)** Structural protein representation of IGHA1, **(B)** IGHA1 and IGHA2 mutations shown in MSA, **(C)** Structural protein representation of IGHA2. The models illustrate the CH1, CH2, and CH3 domains of IGHA1 and IGHA2. In both IgA subclasses, CHS refers to the hydrophilic C-terminal end of secreted IG found in 3’ of the CH3 domain (47). MSA calculated with MUSCLE (v3.8.425), using IMGT/GENE-DB alleles, and visualized in Jalview with BLOSUM62 shading, highlights amino acid substitutions, with novel alleles in bold. Positions of novel mutations are marked in green and known A2m allotypes are marked in orange. These positions are also mapped onto structural models of IgA1 (AF_AFP01876F1) and IgA2 (AF_AFP01877F1), predicted by AlphaFold and visualized in Chimera (48).

In contrast, IGHA2 subclass has three well-defined allotypes: A2m1, A2m2, and A2m3. This study enhances the understanding of IGHA2 allotypes by identifying new alleles. Specifically, IGHA2*06 and IGHA2*07 are classified as A2m1, distinguished by proline residues at positions 115 and 124 in the CH1 domain. Additionally, IGHA2*05, found in Asian sequence IMGT000199, combines characteristics of A2m2 (CH1 S115 and R124) and A2m3 (CH3 Y85.2, E100, M124, I134, and A143), similar to the previously described allele IGHA2*02. This allele, however, is distinguished by two unique mutations: P83S in CH2 and R84Q in CH3, suggesting potential structural variations (Figure 15). The A2m1 allotype is the most prevalent, represented by nine sequences and three distinct alleles across all individuals from different population groups. While the sample size used is limited, these findings contribute to understanding IGHA2 diversity and provide a foundation for larger-scale population studies.

### IGHG1, IGHG2, IGHG3 and IGHG4

For the IgG subclasses, IGHG1, IGHG2, and IGHG3 have defined allotypes while IGHG4 lacks them. Allotypes are subclass-specific amino acid polymorphisms in the constant region that are detectable by antibody reagents. When the variants appear in other subclasses without being polymorphic but remain antibody-detectable; they are termed isoallotypes (8,49). The updated allotype/isoallotype table presented in Figure 16 summarizes the IMGT alleles and their allotype or isoallotype affiliations, along with non-allotypic IGHG variants.

**Figure 16.**
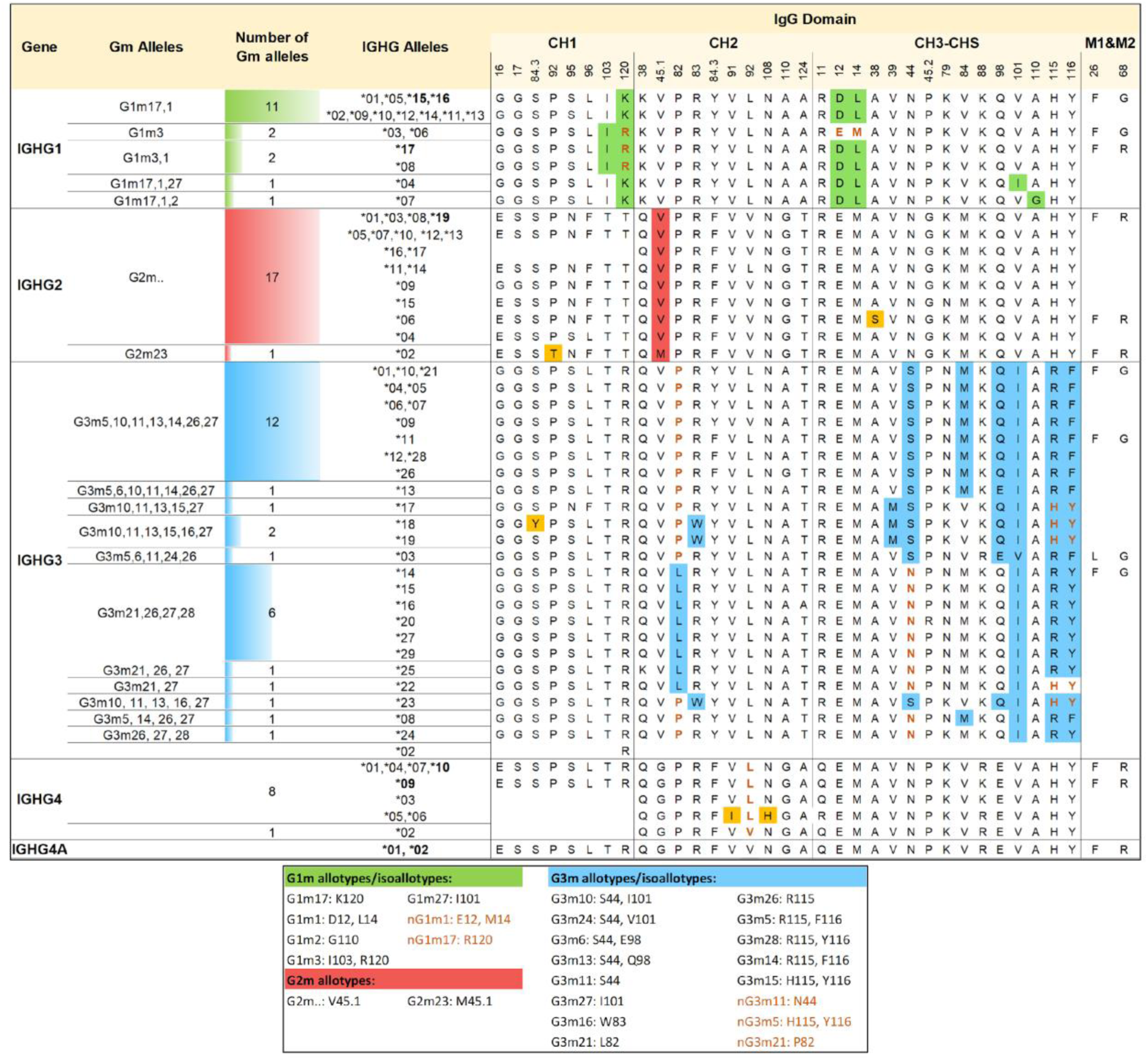
Updated allotype representation for IGHG IMGT annotated genes. This figure presents the correspondence between the Gm alleles and IGHG alleles, highlighting allotypic and non-allotypic variants across exons. With green: G1m allotypes, with red: G2m allotypes, with blue: G3m allotypes, with yellow: unique variants, with orange font: isoallotypes, and with bold: new alleles.

Newly identified IGHG1 alleles, IGHG1*15 and IGHG1*16, combine features from allotypes 1 (D12, L14 at CH3) and 17 (K120 at CH1), and are assigned to the G1m17,1 allele. Similarly, IGHG1*17 belongs to G1m3,1, sharing characteristics from both allotypes 1 and 3 (I103, R120 at CH1). In IGHG2, the newly discovered IGHG2*19 is classified as G2m.. due to V45.1 at CH2. For IGHG4, IGHG4*10 shares the same amino acid sequence as IGHG4*01, IGHG4*04, and IGHG4*07, while IGHG4*09 differs only at position 88 in CH3, where K replaces R, a variation also found in IGHG4*03.

The most prevalent Gm alleles are G1m17,1 for IGHG1 (11 sequences), G2m.. for IGHG2, and G3m5,10,11,13,14,26, and 27, followed by G3m21,26,27,28 (7 sequences) for the IGHG3.

## DISCUSSION

### Assembly Selection Challenges and Integrity: A Key to Reliable IGH Locus Analysis

Assembly selection was pivotal to ensure the integrity and accuracy of the IGH locus analysis. From the 600 assemblies available as of December 1^st^, 2024, 585 were excluded, primarily due to the absence or incompleteness of chromosome 14, or NCBI suppression or diploid status, highlighting the necessity for chromosome-specific resolution in studies of complex genomic loci.

We prioritized assemblies generated by advanced long-read sequencing technologies, such as PacBio Sequel II HiFi and Oxford Nanopore PromethION (50) that offer high resolution and contiguity essential for reconstructing complex loci like IGH. Many were complemented by short-read and scaffolding techniques, including Illumina NovaSeq and Hi-C (51–54). Multimodal assemblies like PGP1 and mHomSap3 exemplify this integrated approach, which enhances phasing accuracy and assembly continuity (51). High-fidelity long reads from PacBio HiFi (>10 kb) and ultra-long reads from Oxford Nanopore (>100 kb) are essential for resolving complex genomic regions, minimizing assembly errors, and enhancing contiguity in repetitive areas (55,56). Assemblies such as T2T-CHM13v2.0 and NA19240 constructed with PacBio Sequel and HiFi sequencing technologies along with advanced pipelines (e.g., T2T, Hifiasm), set a new standard for high-quality reference genomes.

Despite improvements in assembly technology, challenges remain due to the complexity of the locus. We therefore applied IMGT/StatAssembly (25), an IMGT tool dedicated to read-level validation, which confirmed the overall assembly quality and highlighted regions where allele designation was unreliable (Supplementary Figure S2). Further curation was based on IMGT criteria, such as gene order, gap presence and haploid status (57). Assemblies misclassified as haploid (e.g., hg01243.v3.0, NA19240_prelim_3.0), were excluded after confirming their diploid status in the literature (58). When haploid parental assemblies were available, they were retained, validated, and annotated. Assemblies designated as haploid in NCBI were independently verified to mitigate potential discrepancies, for example, PGP1’s haploid status was confirmed through the bioinformatics pipeline with read-based phasing by WhatsHap (59).

Importantly, we chose to retain certain assemblies derived from B lymphocytes despite known limitations due to DJ or VDJ rearrangement. When appropriately annotated, these assemblies offer insight into the dynamic post-rearrangement architecture of the IGH locus in immune cells, offering a biologically relevant view of somatic recombination, gene usage, and structural variability introduced during B-cell maturation (60). However, to ensure accuracy, we did not consider novel alleles located within VDJ rearranged regions, as these may reflect somatic hypermutations rather than germline configurations.

The final selection of 15 assemblies represents a high-quality dataset, providing a robust foundation for further analysis of the IGH locus and human genetic diversity. These assemblies represent a broad range of individuals of diverse population groups, while reference sequences such as T2T-CHM13v20 serve as foundational benchmarks (61). By combining technical validation with an accurate representation of genetic variation, this dataset provides a solid foundation for in-depth exploration of the IGH locus and offers nuanced insights into the complexity of human immunogenetic diversity.

### In-Depth Characterization of Novel Immunoglobulin Allelic Variants: Polymorphisms and IGHC Locus Diversity

#### Expanding the Immunoglobulin Allelic Repertoire: Novel Variants and Implications

Analysis of 15 high-quality and diverse human genome assemblies revealed 120 novel alleles across IGHV, IGHD, IGHJ, and IGHC genes. This substantial number, discovered from a limited sample, emphasizes both the depth of IG gene diversity and the likelihood that many additional alleles remain undiscovered in broader populations. All 120 alleles passed IMGT’s strict quality controls and have been added to IMGT databases, expanding the IMGT® reference directories and the known spectrum of the IGH diversity (16).

The newly identified alleles include functional genes, ORFs, and pseudogenes, each offering further insight into the complexity and evolutionary dynamics of the IGH locus. While functional alleles are central to the adaptive immune diversity, inclusion of ORF and pseudogenes is essential for a comprehensive view of the locus. These sequences reflect the locus’ evolutionary history, including potential past recombination and duplication events, and contribute to understanding its structural complexity (62–64). Pseudogenes, abundant in vertebrates, serve as markers of neutral evolution, making their analysis especially valuable (62,65). Some retain structural features suggesting potential functional or regulatory roles, underscoring their relevance beyond direct antigen receptor diversity (64,66). Their inclusion ensures that reference datasets capture both the functional diversity and the genomic architecture that shapes it.

Overall, these findings, although based on a limited number of individuals from diverse ancestral backgrounds, emphasize the extent of previously unreported immunogenetic variation and underscore the value of high-quality genome assemblies in expanding the known repertoire. The study of the IGH locus diversity enhances our understanding of immune variation and enriches the IMGT reference directories. This enrichment enables more precise analysis of expressed IG repertoires, particularly in patients with autoimmune diseases as part of the IHU Immun4Cure consortium (https://www.immun4cure.fr/). While this enrichment is an initial step, it lays the groundwork for future research into how these variations affect immune response, disease risk, and personalized medicine (67).

### Unveiling Novel Polymorphisms and Variability in the IGHC Genes: Implications for Immunoglobulin Function and Diversity

IGHC gene variability remains underexplored compared to that of IGHV genes, even though it corresponds to an equally complex region, due to multigene deletions and resulting copy number variations (CNVs) observed in healthy individuals (68–70). IGHC genes are crucial in the immune response by determining the isotype of an IG molecule. While IGHG class is well studied for their allotypes and their links to autoimmune, infectious, cancer-related conditions, as well as longevity (71–74), less is known about the other IGHC genes. Interestingly, despite the limited studies, mutations in IGHM have been associated with autosomal recessive agammaglobulinemia (75), and IGHA2 has emerged as a promising breast cancer marker (76).

Our analysis uncovered 31 novel IGHC alleles, primarily within the less-studied IGHD, IGHM, IGHE, and IGHA, suggesting previously underestimated polymorphism levels. Several new alleles include nucleotide mutations in their germline sequences leading to amino acid substitutions potentially affecting the protein structure. For instance, L21F substitution in the CH2 domain of IGHM*05 allele, where a small hydrophobic leucine is replaced by a larger aromatic phenylalanine and the R84.4G substitution in both new IGHD alleles—IGHD*03, and IGHD*05—which involves the replacement of the large and positively charged arginine with the smaller, and flexible glycine, in the DE-TURN of the CH3 domain. Further investigation is essential to determine how these amino acid substitutions influence the IG structure, function, and disease associations.

Furthermore, a substitution detected in the transmembrane region of the IGHE*06 may influence the proteins’ transmembrane domains and potentially impact the receptor activity. These findings are particularly noteworthy, as a recent study emphasized the significance of examining cytoplasmic and transmembrane regions by identifying mutations in this region of the IGHG1 associated with systemic lupus erythematosus (73). Investigating all the exons and transmembrane domains is crucial to fully understand IGHC variations and their disease associations.

### Unveiling the Foundations of Allelic Variation and Diversity Among Individuals from Diverse Ancestries

The human immune system, with its intricate network of genes and complex responses, is shaped by evolutionary forces that have acted on human populations over millennia (77,78). However, the IGH locus, a critical region for the immune function, remains one of the least explored regions, regarding allelic variation observed across individuals of diverse populations.

This study provides a comprehensive annotation of the IGH locus using a selected set of high-quality genome assemblies from individuals of African, European, Saudi, Asian, Puerto Rican and mixed ancestry. Acknowledging the current sample size per group, the analysis reveals substantial allelic variation within this highly polymorphic immune locus.

Despite this variation, some alleles are consistently shared across all six groups, each found in at least one individual per group (55 in total), pointing to a potentially conserved core set of IGH variants essential to human immunity. In parallel, African individual in our cohort exhibits the highest number of unique IGH alleles (72) potentially reflecting Africa’s deep genetic history (79). As Africa is the origin of the greatest human genetic variation, African genome can offer essential context for understanding the full spectrum of immune variation (80,81).

In contrast, European individuals, studied here, display 39 unique alleles, while Saudi, Asian, and mixed, each exhibit up to 22 unique variants (Figure 3), which may reflect population bottlenecks and selective pressures during migration out of Africa (80,82,83), although conclusions are limited by small sample size. The presence of novel alleles unique to individuals of certain populations highlights the extent of variability in the IGH locus and the potential role of local adaptation (Figure 6).

### IGH Locus Novel Insights: Neighboring Alleles, Expression Correlations and Heterozygosity

#### Expression Patterns of Key Genes

While certain genes, such as IGHV1-45, IGHV4-28, and IGHD6-25, have been reported to exhibit low expression across most individuals, raising the possibility that they are non-functional or of limited prevalence (43), our study revealed these three genes to be present in all analyzed individuals (except in rearranged assemblies) and functional. This suggests that their low expression may result from regulatory variation, such as differences in promoter activity and its regulatory elements, or epigenetic modifications, rather than gene absence or loss of function.

#### Correlation of Neighboring Genes in CNV regions

There is a significant correlation between the presence or absence of genes within CNV regions, particularly among functional alleles. Our study confirms previously reported findings and reveals new correlations within the most complex regions of the IGH locus.

The CNV1 indicates that the absence of the IGHV2-70D is associated with the loss of the adjacent genes IGHV1-69-2 and IGHV1-69D (CNV1 form A). Similarly, the CNV2 demonstrates that the absence of the IGHV3-43D corresponds with the absence of the IGHV4-38-2 across the studied assemblies of individuals from diverse population groups (CNV2 form A). A particularly noteworthy observation is seen in CNV3, where the absence of the IGHV4-30-2 and the IGHV4-30-4 (CNV3 forms A, D, G and H) are especially evident in Asian individuals analyzed. Although the allele IGHV3-30-5 is identical to the IGHV3-30, our detailed CNV analysis successfully differentiated these alleles using distance metrics and MSA methods. This approach allowed us to identify genes within the CNV3 even in cases where not all genes were present, effectively resolving a long-standing challenge in this complex IGH region.

In addition, the CNV4 highlights that the duplication of the IGHV3-23 F in a locus, is correlated to the presence of the duplicated genes IGHV(III)-22-2D and IGHV(II)-23-1D. Finally, the CNV5 reveals reciprocal deletions between the genes IGHV3-9 and IGHV1-8, as well as the IGHV5-10-1 and the IGHV3-64D in its two identified forms (forms A and B). Collectively, these findings enhance our understanding of structural variations within the IGH locus and support continued exploration of their potential implications for immune system diversity and functionality.

#### IGH Locus Heterozygosity Across Individuals of Different Ancestral Backgrounds

Our analysis reveals substantial heterozygosity at the IGH locus across the studied individuals, providing insights into allelic diversity. Heterozygosity levels varied, from 11% in the Asian individual to 40% in the African individual, consistent with the potential influence of genetic drift and demographic history. Several IGHV alleles, such as the IGHV1-69, the IGHV3-53, the IGHV3-48, the IGHV3-49, and the IGHV3-11, exhibited heterozygosity in over 50% of analyzed individuals, consistent with their known polymorphic nature (43). Notably, the IGHV1-69 showed distinct maternal-paternal allele combinations in some individuals (*12/*14 in African, *01/*06 in mixed and *12/*01 in Puerto Rican) whereas others (Asian, Saudi), were homozygous, illustrating allele diversity within this locus (Figure 5). Consistent with previous findings, the IGHJ6 was frequently heterozygous, with about 30% heterozygosity, often involving *02 and *03 alleles, like the Eastern European Ashkenazi Jewish individual (43). Although the limited dataset size, the heterozygosity observed in this study could help our efforts to understand the IGH allelic variation relevant to personalized immune repertoires.

### Exploring the CNV Variability of the IGH Locus: Potential Population-Specific Implications for Immune Function and Disease

CNVs significantly contribute to genomic variation, impacting both phenotypic traits and disease susceptibility. Our investigation into the different IMGT CNV forms has revealed distinct configurations across individuals. For example, in CNV4, the African, Puerto Rican, Asian, and Saudi individuals predominantly share two distinct forms, the regional genetic diversity that underpins these groups. The African individual, notably, uniquely carries the CNV2 form B, highlighting its potential genetic divergence.

In the mixed individual, we observed distinct variability between maternal and paternal IMGT CNV forms. For instance, CNV5 appeared as form A in the maternal assembly and form B in the paternal assembly, demonstrating differences in gene content between IGHV3-11 and IGHV3-7, although both forms retained two functional genes within this region. Notably, the CNV4 and CNV6 displayed similar forms in both maternal and paternal assemblies, indicating a more consistent configuration.

The CNV3 represents the most complex segment of the IGH locus, encompassing both functional genes, alongside several pseudogenes, and displaying significant variability, and a critical role in the dynamic genetic landscape of the IGH locus. Both the Asian individual and the maternal assembly of the Saudi individual show the fewest genes, with only one functional gene (the IGHV3-33 for the Saudi and the Asian paternal assembly and the IGHV3-30-3 for maternal assembly). In contrast, the most comprehensive CNV3 form C includes all identified genes, is shared among the European, Puerto Rican, and mixed ancestries, indicating potential common evolutionary patterns. Despite overall allele variability, certain alleles, such as the IGHV3-30*18 F, the IGHV3-33*01 F, and the IGHV4-31*03 F, are consistently conserved across diverse ancestries, with rare exceptions like the IGHV3-33*06 F in the paternal African Yoruba assembly. This conservation suggests potential evolutionary advantages crucial for immune function. Understanding the functional implications of conserved alleles like IGHV3-30*18 might be crucial for developing immunotherapies and vaccines, to diverse populations. Our examination of the CNV7 forms, across individuals from different ancestries, highlights remarkable variability in the IGHC genes. Notably, assemblies from mixed ancestry, and those from the Asian individual consistently feature all 11 IGHC genes. In contrast, the Eastern European Ashkenazic Jewish assembly reveals a more complex configuration, encompassing all 12 IGHC genes, including the novel IGHG4A gene.

This new discovery sheds light into the current limited knowledge regarding the prevalence of this IGHG4 duplication. IGHG4A, identified in 31.25% of the studied sequences, aligned with previous studies (40-44%), and is distinct from IGHG4 due to an amino acid substitution, V92L (84,85). Across all duplication cases, the IGHG4 retained the L92 mutation, whereas the IGHG4A exhibited the V92 variant, consistent with prior findings, while the rare combination of IGHG4A L92 and IGHG4 L92, was not observed in our dataset (84). Regarding non-duplicated IGHG4 sequences, the L92 variant in 56.25% of cases closely matches the reported prevalence of 55.6%, suggesting consistency with broader population patterns. On the contrary, the IGHG4 with the V92L in 12.5% of the cases, was significantly higher than the 2.2% reported (84).

These insights into IGHC gene variability are crucial given the significant role of antibody isotype diversity in modulating immune responses across diverse disease contexts. Specifically, IgG4, though the least abundant IgG subclass in serum, is uniquely linked to both immune tolerance and chronic inflammation (86). Furthermore, elevated IgG4 responses have been observed in multiple cancers (87–89), with a recent study showing that IgG4 can inhibit anti-tumor immunity (90). Increased IgG4 levels are also characteristic of chronic inflammatory diseases, including rheumatoid arthritis and IgG4-related disease, highlighting its role in immune regulation (91,92). The identified duplication of the IGHG4 (IGHG4A) may lead to increased IgG4 expression, and enhance its immunomodulatory effects, potentially influencing immune responses in chronic inflammation and tumor immunity, representing a promising area for future research.

### The Foundations of the IMGT haplotype Database

The combination of the variable CNV forms across the loci of different individuals, shapes their IMGT haplotype. By identifying eight distinct IMGT haplotypes, this analysis establishes a foundation for the development of an IMGT haplotype database with the goal of identifying profiles potentially relevant to cancer and autoimmune diseases.

## CONCLUSION

This study provides an in-depth characterization of the IGH locus, a critical component of the genome involved in adaptive immunity. By analyzing 15 high-quality genome assemblies from individuals of diverse ancestral backgrounds, we identified substantial genetic variation across the locus. This includes 124 novel IGH alleles spanning the variable, diversity, joining, and constant genes, substantially enriching the IMGT reference database. We highlight significant genetic diversity among individuals and illustrate complex patterns of CNVs through distinct CNV forms, reflecting the complexity of the region’s structural variation. The identification of a novel gene, IGHG4A, along with previously unreported amino acid substitutions in IGHC alleles points to previously underestimated polymorphisms with potential implications for immunoglobulin structure and function. This research establishes a crucial foundation for the emerging IMGT® haplotype database, which will be pivotal for supporting future studies in population-specific immune profiles, autoimmune susceptibility, and the development of personalized immunogenomics in various disease contexts.

## DATA AVAILABILITY

IMGT^®^ is freely available online for academics and non-profit use at http://www.imgt.org/. All the databases and tools referred to in this article are accessible from the IMGT® webpage. Several of the annotated sequences are available on the Third Party Annotation (TPA) database of NCBI (BK063800.1, BK063801.1, BK063802.1, BK068298.1, BK068299.1, BK070981.1 and BK070982.1). The data underlying this article are available in the article and in its online supplementary material.

## SUPPLEMENTARY DATA

Supplementary Data are available at NAR Online.

## ACKNOWLEDGMENTS

We sincerely appreciate the IMGT® team for their dedication and steadfast enthusiasm, and we particularly acknowledge Dr. Taciana Manso for her valuable comments in our study. Over the course of this approximately five-year project, several Erasmus+ and French university internship students participated in IMGT biocuration training, most for a duration of two months. We appreciate the energy and fresh perspectives they bring to the team and we list them in alphabetical order: Kawtar Daky, Eleni Kaloudi, Vasiliki Panagiotou, Amélie Torino, and Michaela Margarita Triantafyllou. We would also like to thank Dr. Anthony Boureux, for introducing us to the CountTags tool, which significantly contributed to our work.

## AUTHOR CONTRIBUTIONS

Conceptualization, Methodology, Validation, and writing – review & editing: A.P., M.G., G.F., J.J.-M., G.Z., P.D., V.G., and S.K.; Investigation, Data Curation, Visualization and Writing – original draft: M.G., and A.P.; Supervision: V.G. and S.K. Funding acquisition: S.K. All authors have read and agreed to the published version of the manuscript.

## FUNDING

IMGT® is currently supported by the Centre National de la Recherche Scientifique (CNRS) and the University of Montpellier. IMGT® is a member of the French Infrastructure ‘Institut Français de Bioinformatique’, IFB as well as member of BioCampus, MAbImprove and IBiSA. This work was granted access to the High Performance Computing (HPC) resources of Meso@LR and of Centre Informatique National de l’Enseignement Supérieur (CINES), to Très Grand Centre de Calcul (TGCC) of the Commissariat à l’Energie Atomique et aux Energies Alternatives (CEA), and to Institut du développement et des ressources en informatique scientifique (IDRIS) under the allocation 036029 (2010–2025) made by GENCI (Grand Equipement National de Calcul Intensif). We acknowledge the support of Immun4Cure University Hospital Institute ‘Institute for innovative immunotherapies in autoimmune diseases’ (France 2030 / ANR-23-IHUA-0009) and also by the Key Collaborative Research Program of the ‘Alliance of International Science Organizations’ (ANSO-CR-KP-2022-09). SK acknowledges the financial support of the IUF to IMGT.

## CONFLICT OF INTEREST

The authors declare that they do not have any conflict of interest for the work carried out in this manuscript.

## Notes

### Competing Interest Statement

The authors have declared no competing interest.

