## Supplementary Figure S1 for "IMGT® Analysis of the Human IGH Locus: Unveiling Novel Polymorphisms and Copy Number Variations in Genome Assemblies from Diverse Ancestral Backgrounds"

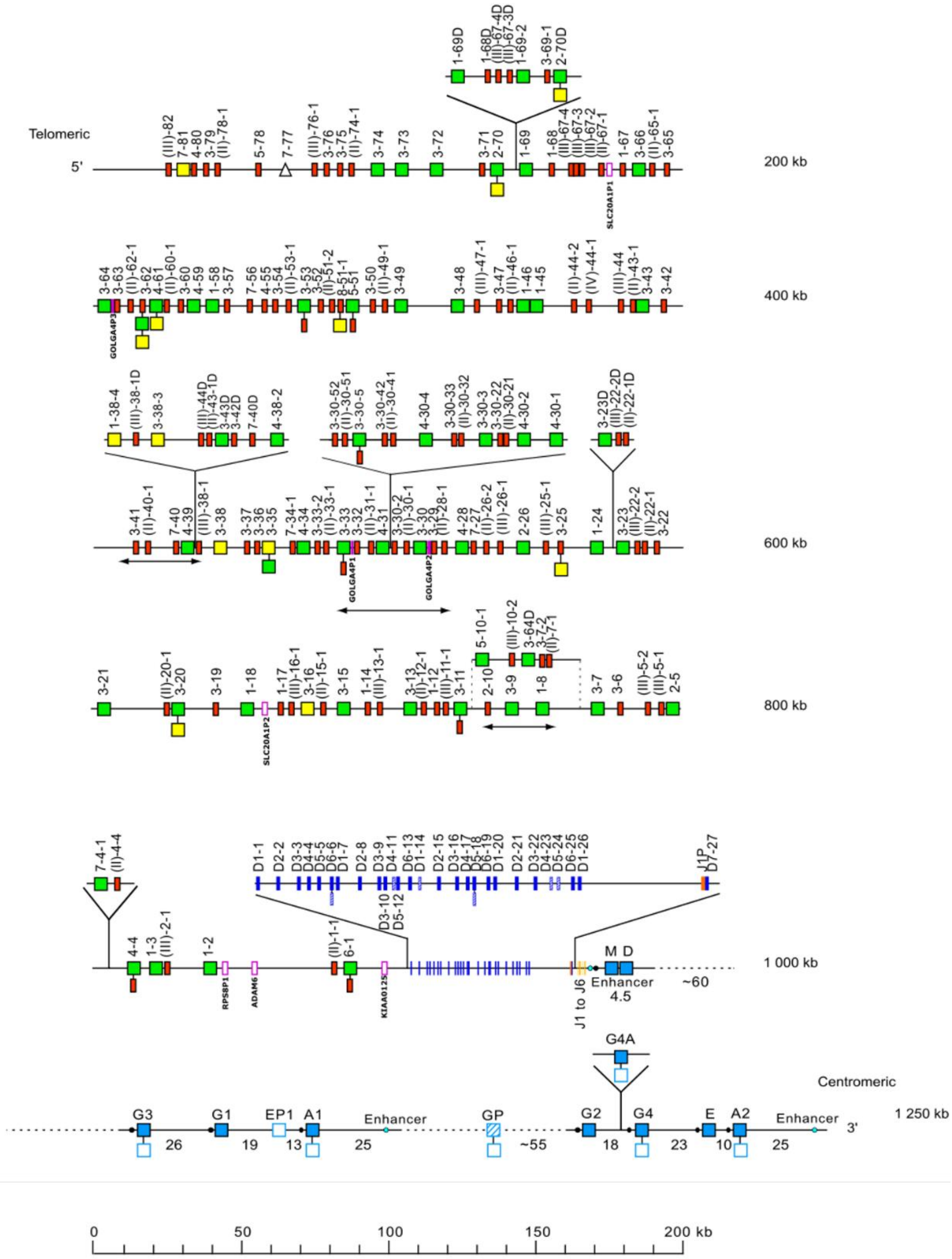


**Supplementary Figure S1.** Holistic IMGT® schematic representation of the human IGH locus. This figure provides a comprehensive overview of the organization of the human IGH locus, reflecting the functional and structural complexity of the locus. Located on chromosome 14 (14q32.33), the locus comprises IGHV, IGHD, IGHJ, and IGHC genes. Each gene is depicted as a box, with its color indicating functionality and gene type, consistent with the IMGT® color menu (https://www.imgt.org/IMGTScientificChart/RepresentationRules/colormenu.php#LOCUS). Genes shown with two boxes indicate cases where the gene has been identified with different functionalities in various alleles. Double arrows indicate insertion/deletion of genes, indicating common Copy Number Variations within the locus. This representation is continuously updated on the IMGT® webpage (https://www.imgt.org/IMGTrepertoire/index.php?section=LocusGenes&repertoire=locus&species=human&group=IGH&assembly=Holistic_IMGT_reference) as new genes or new alleles functionalities are identified.
