## Supplementary Figure S2 for "IMGT® Analysis of the Human IGH Locus: Unveiling Novel Polymorphisms and Copy Number Variations in Genome Assemblies from Diverse Ancestral Backgrounds"

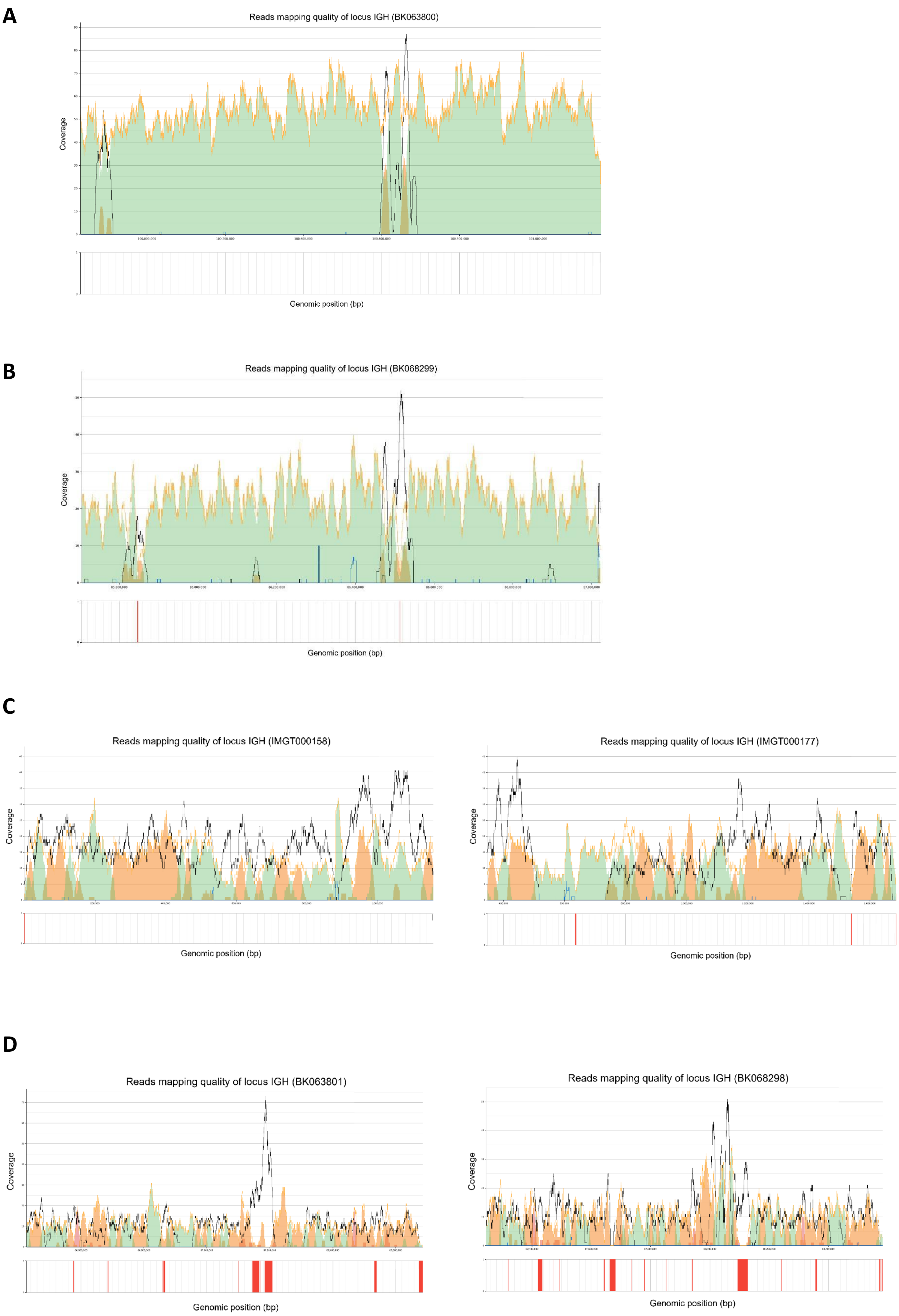
A
Reads mapping quality of locus IGH (BK063800)
Coverage
Genomic position (bp)
B
Reads mapping quality of locus IGH (BK068299)
Coverage
Genomic position (bp)
C
Reads mapping quality of locus IGH (IMGT000158)
Coverage
Genomic position (bp)
Reads mapping quality of locus IGH (IMGT000177)
Coverage
Genomic position (bp)
D
Reads mapping quality of locus IGH (BK063801)
Coverage
Genomic position (bp)
Reads mapping quality of locus IGH (BK068298)
Coverage
Genomic position (bp)

**E**

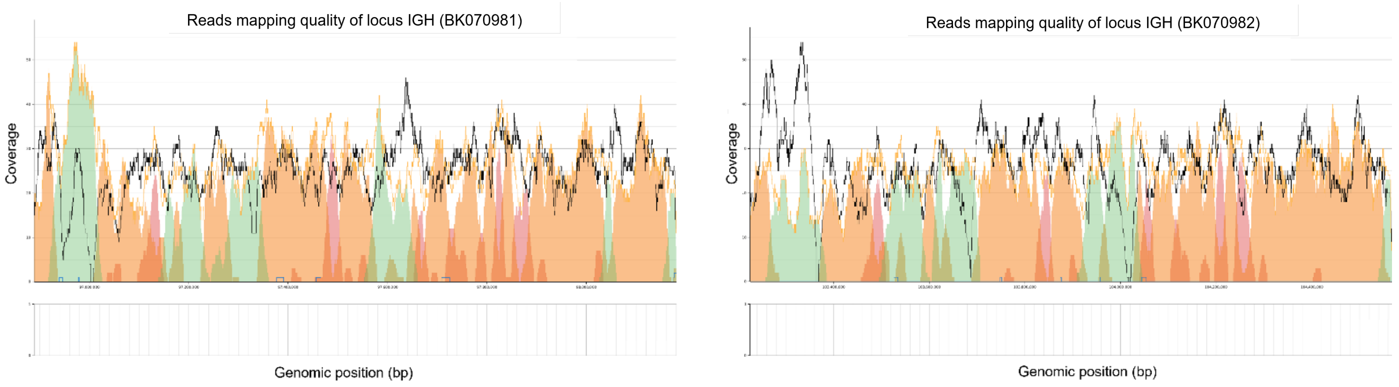
Reads mapping quality of locus IGH (BK070981)
Reads mapping quality of locus IGH (BK070982)
Coverage
Genomic position (bp)

**F**

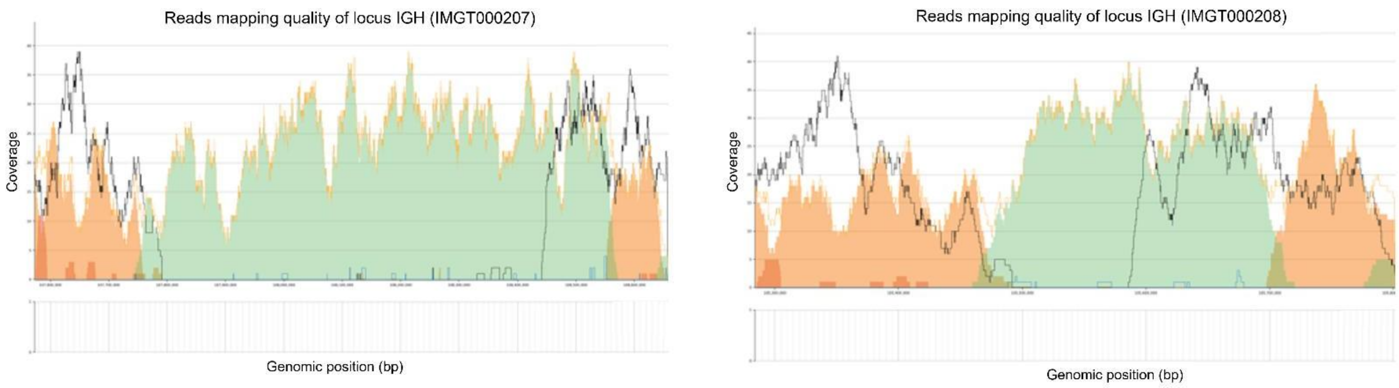
Reads mapping quality of locus IGH (IMGT000207)
Reads mapping quality of locus IGH (IMGT000208)
Coverage
Genomic position (bp)

**G**

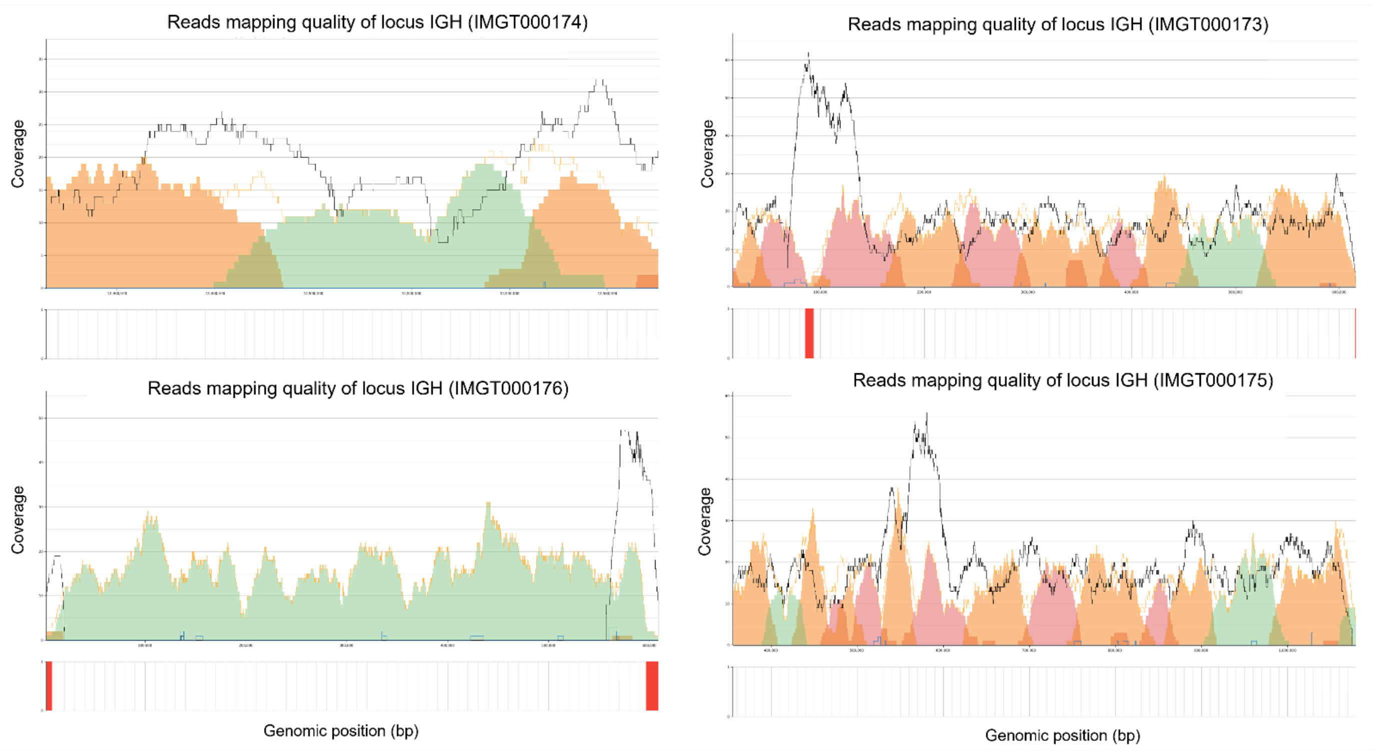
Reads mapping quality of locus IGH (IMGT000174)
Reads mapping quality of locus IGH (IMGT000173)
Reads mapping quality of locus IGH (IMGT000176)
Reads mapping quality of locus IGH (IMGT000175)
Coverage
Genomic position (bp)

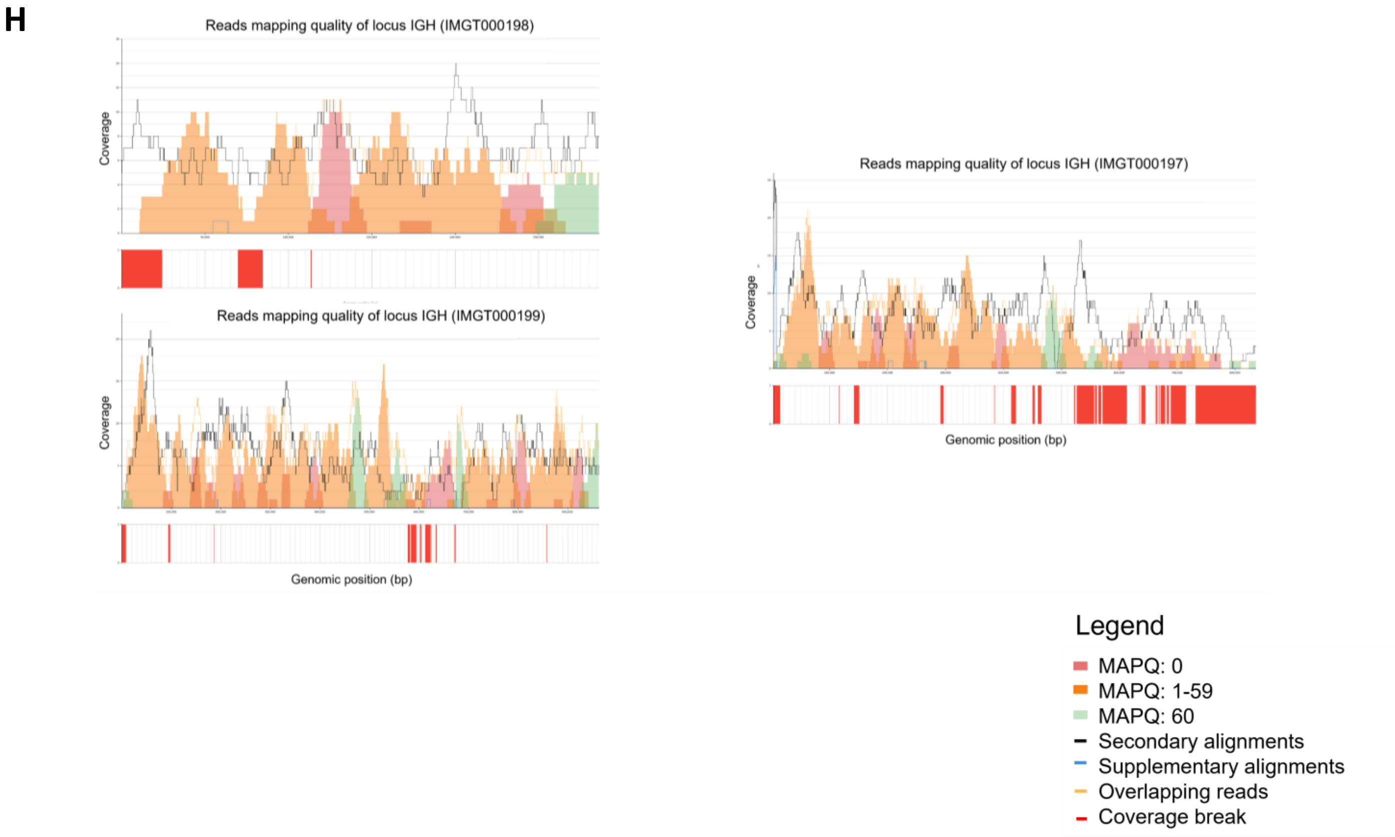


**Supplementary Figure S2.** The graphs depict the mapping quality of reads within the IGH locus, extracted from the corresponding assemblies. They were generated using IMGT/StatAssembly (version v0.1.4), which analyses the BAM alignments derived from the mapping of HiFi reads to the associated assemblies. Green represents regions with Phred quality score 60, orange indicates Phred score between 1-59, and red denotes a score of 0. Black highlights secondary alignments, while the red bars represent coverage breaks (default ≤3). Mapping quality of IGH locus for each studied assembly: (A) Assembly T2T-CHM13v2.0 (BK063800) (B) Assembly PGP1v1 (BK068299) (C) Maternal and Paternal assembly of NA19240 (NA19240.pri.mat.f1_v2 -IMGT000158, NA19240.alt.pat.f1_v2 - IMGT000177) (D) Maternal and Paternal assembly of mHomSap3 (mHomSap3.mat-BK063801, mHomSap3.pat- BK068298) (E) Maternal and Paternal assembly of KSA001 (ASM3717763v1-BK070981, ASM3717755v1- BK070982) (F) Maternal and Paternal assembly of NA24385 (hg002v1.0.1.mat- IMGT000207, hg002v1.0.1.pat - IMGT000208) (G) Maternal and Paternal assembly of HG01243 (HG01243.pri.mat.f1_v2 - IMGT000175, IMGT000176, HG01243.alt.pat.f1_v2 - IMGT000173, IMGT000174) (H) Maternal and Paternal assembly of NA24631 (HG005.pri.mat.f1_v2 - IMGT000197, IMGT000198, HG005.alt.pat.f1_v2 - IMGT000199).
