## Supplementary Figure S3 for "IMGT® Analysis of the Human IGH Locus: Unveiling Novel Polymorphisms and Copy Number Variations in Genome Assemblies from Diverse Ancestral Backgrounds"

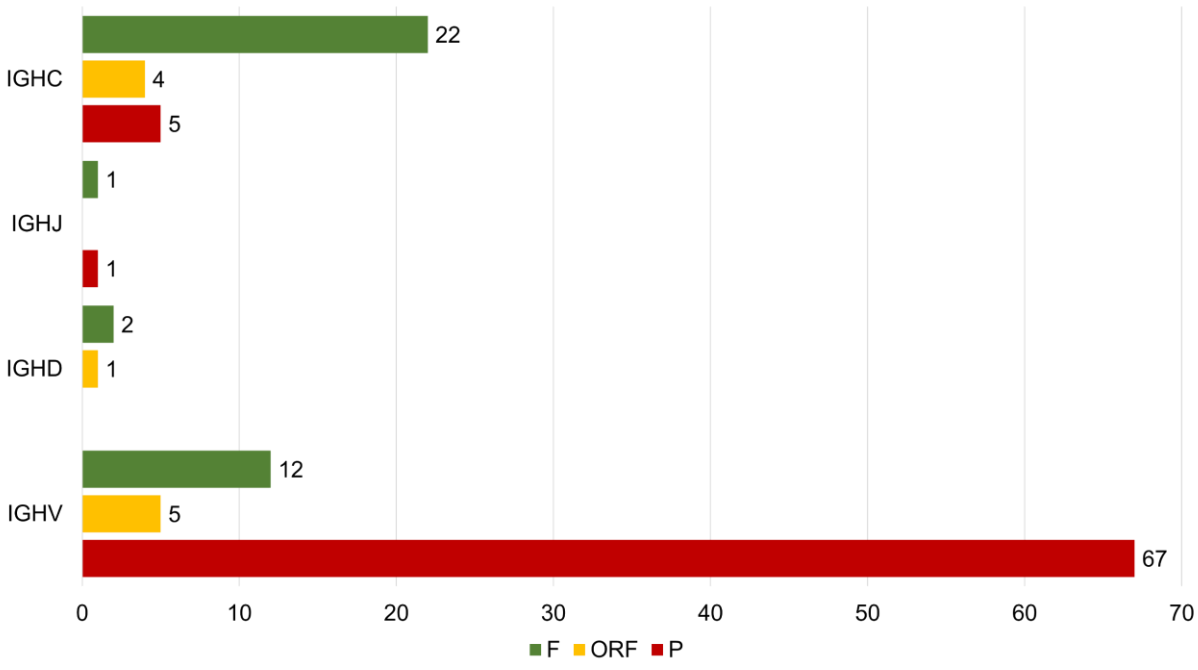


**Supplementary Figure S3.** Distribution of novel alleles per functionality (F, ORF, P) and gene type (IGHV, IGHD, IGHJ, IGHC).
